# THE DUAL-TASK COST IS DUE TO NEURAL INTERFERENCES DISRUPTING THE OPTIMAL SPATIO-TEMPORAL DYNAMICS OF THE COMPETING TASKS

**DOI:** 10.1101/2020.12.07.414953

**Authors:** Diego Mac-Auliffe, Benoit Chatard, Mathilde Petton, Anne-Claire Croizé, Florian Sipp, Benjamin Bontemps, Adrien Gannerie, Olivier Bertrand, Sylvain Rheims, Philippe Kahane, Jean-Philippe Lachaux

**Author notes:** DYCOG address: **Centre de Recherche en Neurosciences de Lyon,** CRNL Inserm U1028 - CNRS UMR5292 - UCBLyon1, CH Le Vinatier - Bâtiment 452, 95 bd Pinel, 69675 Bron Cedex, Rhône-Alpes, France, DYCOG /53.

## Abstract

Dual-tasking is extremely prominent nowadays, despite ample evidence that it comes with a performance cost: the Dual-Task (DT) cost. Neuroimaging studies have established that tasks are more likely to interfere if they rely on common brain regions, but the precise neural origin of the DT cost has proven elusive so far, mostly because fMRI does not record neural activity directly and cannot reveal the key effect of timing, and how the spatio-temporal neural dynamics of the tasks coincide.

Recently, DT electrophysiological studies in monkeys have recorded neural populations shared by the two tasks with millisecond precision to provide a much finer understanding of the origin of the DT cost. We used a similar approach in humans, with intracranial EEG, to assess the neural origin of the DT cost in a particularly challenging naturalistic paradigm which required accurate motor responses to frequent visual stimuli (task T1) and the retrieval of information from long-term memory (task T2), as when answering passengers’ questions while driving.

We found that T2 elicited neuroelectric interferences in the gamma-band (>40 Hz), in key regions of the T1 network including the Multiple Demand Network. They reproduced the effect of disruptive electrocortical stimulations to create a situation of dynamical incompatibility, which might explain the DT cost. Yet, participants were able to flexibly adapt their strategy to minimize interference, and most surprisingly, reduce the reliance of T1 on key regions of the executive control network – the anterior insula and the dorsal anterior cingulate cortex – with no performance decrement.

**HIGHLIGHTS:**

- First direct evidence in humans of neural interferences between two tasks.
- First explanation of the Dual-Task cost at the neural level in humans.
- First Dual-Tasking study with intracranial EEG in naturalistic conditions.

## INTRODUCTION

With increased time-pressure and easy access to digital entertainment, there is a growing temptation to do more in less time, and unsurprisingly, multi-tasking has become a dominant feature of our busy lives, often heralded as a means of living life to its fullest. Yet, years of fundamental and applied psychology research have firmly established that Dual-Tasking (DT) leads to a decrement in performance, called the DT interference effect or DT cost : more errors and slower responses (Welford, 1952; Pashler, 1994; Huestegge et al, 2014). Unsurprisingly, this emerging societal trend has raised serious concerns in various fields, ranging from education to road safety (Strayer & Drews, 2007; Zheng et al, 2014, Pashler et al, 2013).

Our understanding of the DT cost has grown remarkably since the 50’s, with two major lines of explanation to date (Marois & Ivanoff, 2005; Koch et al, 2018; Strobach et al, 2018): “central bottleneck” theories postulate that all tasks that involve flexible stimulus-response associations rely on a central response selection component that can only process one task at a given moment, thereby imposing a waiting time to the other task (Welford, 1952; Pashler, 1994). In contrast, “resource-sharing” theories suggest that two tasks can run in parallel, but compete for limited cognitive resources, which leads to performance decrement (Kahneman, 1973; Logan & Gordon, 2001; Meyer & Kieras, 1997; Miller et al, 2009; Navon & Miller, 2002; Tombu & Jolicoeur, 2003; Wickens, 1984). Over the years, both theoretical frameworks have led to an arborescence of formal models refining many aspects of that global picture, but the main distinction is still prevalent today.

Prominent research groups have argued that the two theories lead to similar predictions at the behavioral level (Byrne & Anderson, 2001; Navon & Miller, 2002; Tombu & Jolicoeur, 2003), and many investigators turned to neuroimaging to search for evidence for bottlenecks and shared resources in fMRI data (e.g. Worringer et al, 2019 for review). A typical result – compatible with both theories – is that tasks that activate common brain regions in a single-task condition are more likely to interfere, in line with the overlap hypothesis formulated at the turn of the 1980’s (Kinsbourne & Hicks, 1978; Roland, 1985; Klingberg & Roland, 1997). In addition, several studies have reported results supporting more specifically one of the two scenarios, but with no emerging consensus so far (Nijboer et al, 2014, Watanabe & Funahashi, 2018; Worringer et al, 2019). Some studies used time-resolved fMRI (e.g. Tombu et al, 2011) to show that the peak of the BOLD activation associated with a response-selection (RS) component could be delayed by a secondary task performed at the same time, in support of the bottleneck theory. Yet that effect could also be explained by a longer processing time of a RS component used by both tasks in parallel. As an example, Just and colleagues compared the BOLD signal in the DT and ST conditions in voxels commonly activated by two tasks, and found that the DT activity was often less than predicted if ST activations simply added up (Just et al, 2008). This was taken as a strong indication that tasks indeed compete for limited resources, for instance via a time-sharing mechanism (Nijboer et al, 2014), provided a linear relationship between the BOLD signal and the amount of resources devoted to a task. Such lack of convergence might be due to two intrinsic limitations of fMRI: its poor time-resolution – even with time-resolved fMRI – and the fact that neural activity is measured indirectly. For instance, it is often implicitly assumed that a weaker BOLD response corresponds to a decrease of the resources allocated to a task and poor behavioral performance.

Yet it can also be caused by a shorter processing time during efficient trials with fast reaction times. fMRI lacks the temporal precision to distinguish between stronger and longer responses, which might have opposite behavioral correlates. It is also too slow to reveal fast time-sharing mechanisms proposed by some proponents of the resource theories or it cannot measure accurately the onset of neural processes and detect the waiting periods and queuing effects predicted by bottleneck theories (see Watanabe & Funahashi, 2018 for a review of the limits of fMRI to study DT). In fact, if a bottleneck is a specific neuronal population that is required by the two tasks at the exact same time, it can only be identified with focal recordings of that population with a very high temporal precision.

For those reasons, there is a growing sense that fMRI and behavioral measurements alone cannot provide a full explanation of the DT cost. Watanabe & Funahashi recently called for a new wave of DT studies using direct electrophysiological recordings in animals (Watanabe & Funahashi, 2018). With their unlimited temporal precision and direct measure of the neural activity, electrophysiological recordings can reveal the neural substrates of individual cognitive components of a task, in real-time, and detect interferences produced by a secondary task. This might prove essential for the study of naturalistic DT conditions in which the timing of the two tasks relative to each other is not precisely controlled, as direct neural recordings can reveal if specific components of the two tasks operate at the same time.

Yet, animal studies often include an extensive training period, which can affect DT abilities (e.g. Ophir et al, 2009; Schubert et al, 2015) and there are obviously ill suited to investigate high-level cognitive functions specific to humans. However, precise electrophysiological recordings are also performed in humans with intracranial EEG (iEEG). iEEG has been used for decades to study the large-scale spatio-temporal organization of cognition (Lachaux et al, 2012), and we recently used that technique to describe with millimetric and millisecond precision the global neural dynamics of a continuous attention task, which monitors in real-time the ability of participants to stay-on-task (BLAST) (Petton et al, 2019). As BLAST requires participants to process fast repeating visual stimuli to produce accurate motor responses, we reasoned that the task could be combined with a memory-retrieval task to investigate the neural mechanisms of DT interference in situations that require attention to be split between external and internal information. This division is found in many real-life DT situations, for instance when a driver searches for an answer to a passenger’s question.

To maximize ecological validity, we recorded untrained human participants as they performed BLAST is isolation, or combined with a self-paced Verbal Fluency Task, in direct interaction with an experimenter. We report for the first time in humans how two tasks interfere at the local neural level, through incompatible neural dynamics in cortical regions supporting language, visual perception and cognitive control. We show that the secondary task can bend the neural dynamics of the primary task away from its optimal spatio-temporal pattern, through ‘neuroelectric’ interferences, but that radical strategy changes can also minimize that effect.

## MATERIALS AND METHODS

### Participants

Intracranial EEG recordings (iEEG) were collected in twelve patients candidates for epilepsy surgery at the Epilepsy Departments of the Grenoble and Lyon Neurological Hospitals. Eleven to fifteen semi-rigid, multi-lead electrodes were stereotactically implanted in each patient (stereotactic EEG — SEEG, a special type of iEEG) (Kahane et al, 2003). Each electrode had a diameter of 0.8 mm and, depending on the target structure, consisted of 10 to 15 contact leads 2 mm wide and 1.5 mm apart (i.e. 3.5 mm center-to-center, DIXI Medical Instruments). Selection of sites to implant was entirely based on clinical purposes, with no reference to the present experimental protocol. All electrodes showing traces of epileptiform activity were excluded from the present study (visual inspection by the clinical team). All participants were native French speakers with normal or corrected-to-normal vision, and had given their written informed consent; all experimental procedures were approved by the Institutional Review Board and by the National French Science Ethical Committee.

### Stimuli and Tasks

#### BLAST (T1)

The continuous attention task was adapted from the BLAST paradigm published by our group (Petton et al, 2019) to detect transient failures of executive attention. In short, BLAST repeatedly asks participants to find a target letter (the Target) in a subsequent two-by-two square array of four letters (the Array), with new letters every trial (Target and Array) (Fig.1B). Each trial starts with the central presentation of the Target for 200 ms, followed by a mask (# sign) for 500 ms, until the presentation of the Array which remains on screen until the manual response, or for a maximal duration of 3000 ms. The next trial starts after a 800 ms pause, with no visual or auditory feedback of any kind.

**FIGURE 1.**
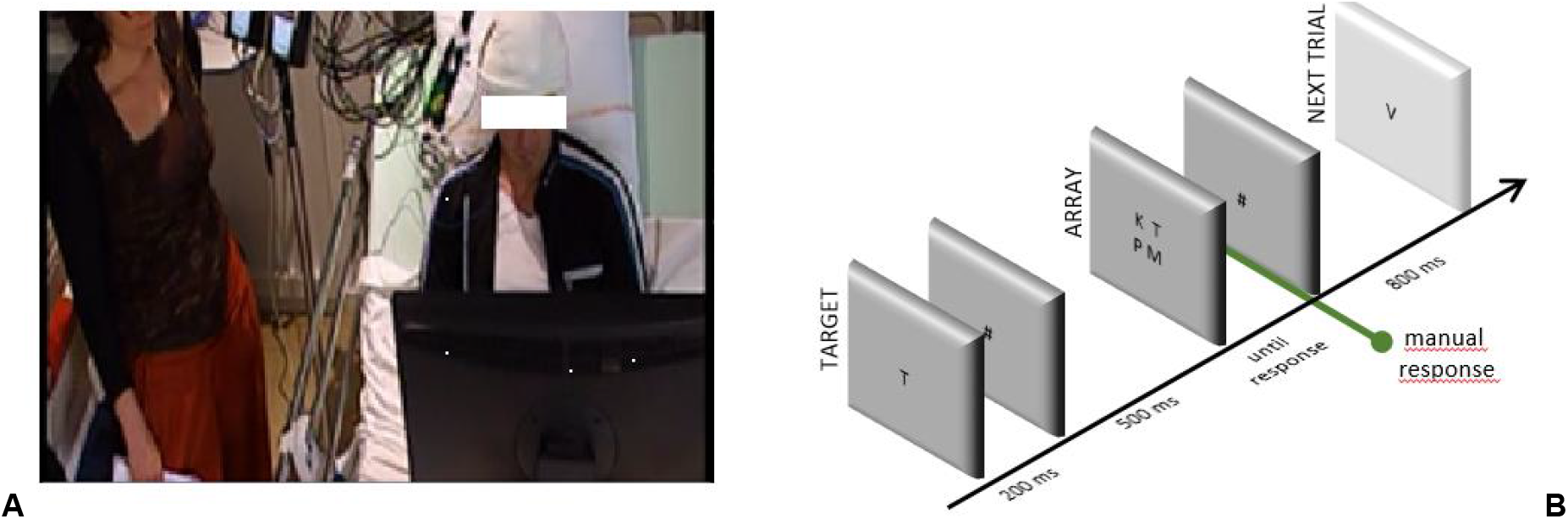
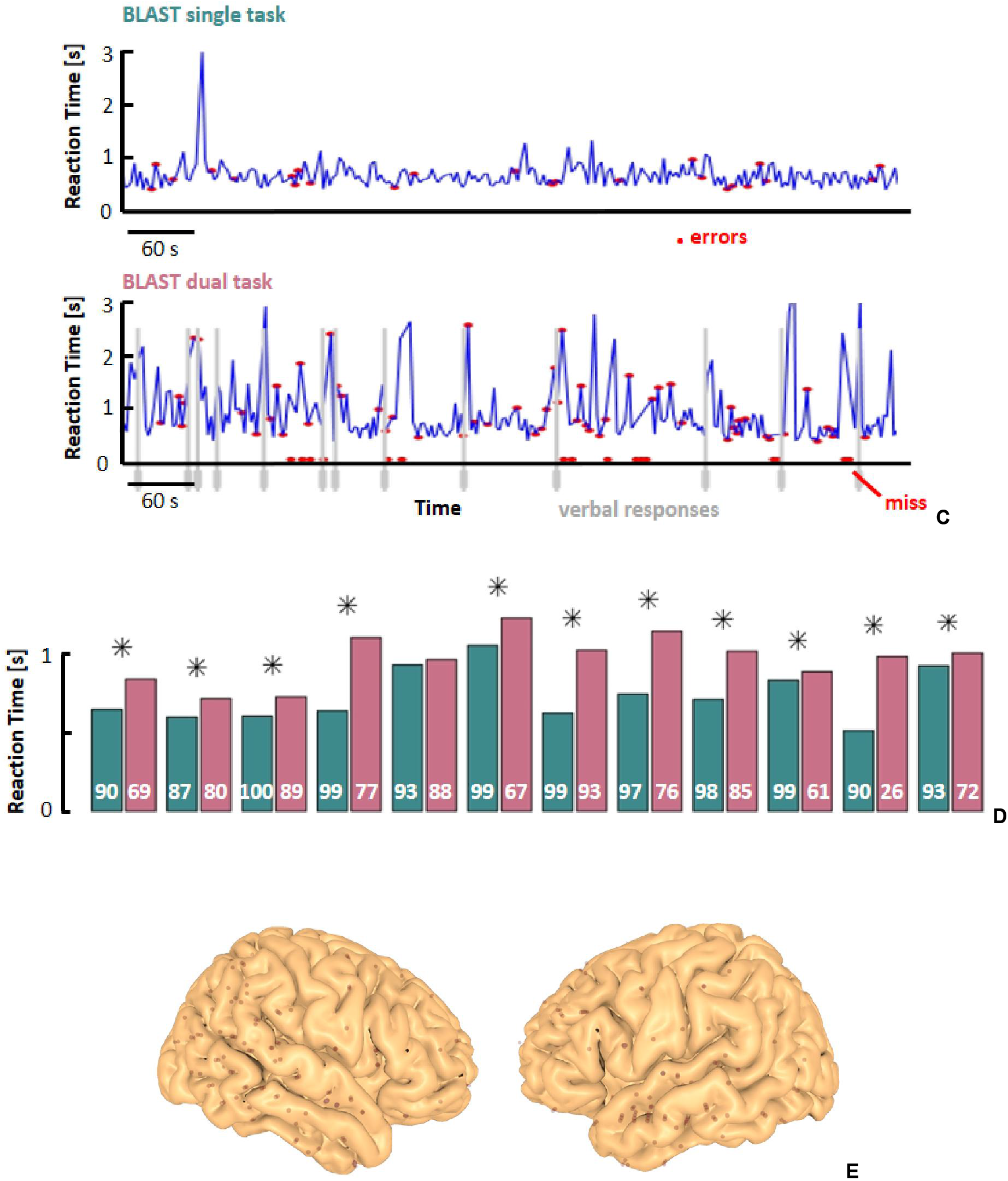
Experimental design. A) Dual-Task set-up: the experimenter is standing or sitting at the bedside, in the peripheral visual field of the participant while the participant is performing BLAST (T1) — the verbal interaction is limited to a strict minimum, necessary for the VFT (T2). B) Schematic depiction of T1 (see methods for details). C) Example of behavioral performance during T1, in the Single-Task (ST) and Dual-Task (DT) conditions. The interference between the two tasks is clearly visible with slower and less accurate responses in the DT condition (vertical gray lines indicate VFT overt responses). D) Mean reaction time during T1 for all participants, in the ST (teal) and DT (magenta) conditions (a star sign indicates a significant difference between the two conditions, two-sample t-test; p<0.05). Accuracy is indicated in white (percentage of correct responses). E) Entry points of iEEG depth-electrodes across all participants, projected onto the MNI-single subject brain template.

Stimuli were delivered on a PC using the Presentation^®^ software (Version 18.0, Neurobehavioral Systems, Inc., Berkeley, CA), synchronized with the EEG acquisition system. The letters were presented foveally in black on a light gray background. Participants pressed a button on a gamepad with their preferred (resp. non-preferred) index finger to indicate if the Target was absent (resp. present) in the Array. The trials sequence was pseudorandomized, with an equal number of target-present and target-absent trials. Performance was measured over a total of 250 trials, for a total duration around ten minutes (depending on reaction times).

The global instruction was to avoid errors and to keep a steady and reasonably fast pace, with an explicit analogy to car-drivers who avoid accidents at all costs, but move forward at a decent speed. BLAST was first performed by participants in a single-task (ST) condition, with no exogenous distractions, and then in a dual-task (DT) condition, simultaneously with a Verbal Fluency Task (Borkowski et al, 1967) (see below).

#### Verbal Fluency Task (T2)

The Verbal Fluency Test (VFT) is primarily used in clinical settings as a diagnostic tool (Andreou & Trott, 2013), or to assess cognitive functions in patients suffering from Alzheimer Disease or schizophrenia (Sumiyoshi et al, 2001; Duff-Canning et al, 2004; Woods et al, 2016). It provides a good evaluation of the ability to retrieve lexical knowledge (Federmeier et al, 2002, 2010) and more generally, of executive control (Fitzpatrick et al, 2013; Shao et al, 2014). In its most typical versions, participants are asked to provide as many words of a given semantic category (semantic VFT) or starting with a given letter (letter or phonemic VFT), as possible in a given time (Shao et al, 2014). VFT performance relies heavily on language and executive processes (Weiss et al, 2003) and at the neural level, on the left Inferior Frontal Gyrus (IFG, including Broca’s area), the dorsolateral Prefrontal Cortex (PFC), the premotor cortex, and the right cerebellum (McGraw et al, 2001).

To maximize the demand on the central executive system, we combined the two VFT variants into one task: a letter was chosen at the beginning of the experiment and participants had to provide names of a given category (animals, names of people, towns, tools …) starting with that letter. Category names were given by an experimenter sitting in the same room as the participant, in its remote peripheral field (Fig. 1A). The timing of the VFT was not computerized so that the switch to a new category occurred as soon as the participant showed obvious signs s/he was running out of answers —, which was better appreciated by an experimenter than by a software. Apart from providing new category names, the experimenter remained still and silent throughout the experiment. VFT performance was analyzed offline from a video recording of the session. To mimic naturalistic conditions, participants received no instruction to prioritize BLAST or the VFT.

In addition, this study makes use of several datasets recorded in separate sessions while participants performed short functional localizers designed to identify iEEG sites involved in visual perception, auditory perception, semantic and phonological processing, verbal and visuo-spatial working memory as well as visual attention respectively (Vidal et al, 2011; Ossandón et al, 2012). Although those data cannot be fully described in this focused paper, we will refer to them each time they provide important insights regarding the function supported by the cortical sites of interest for our study.

### Behavioral Analysis

Our DT design came with two advantages: a) the experimental situation resembled naturalistic conditions (when asked a question that requires an access to long-term memory, people must rarely give their answer within a precise, computerized time-limit, besides TV shows and video games) (Tikka & Kaipainen, 2014), and b) it let participants choose the optimal time-windows to carry out the secondary VFT task T2. The downside was that it was difficult to assess precisely when participants engaged in T2. The only periods when participants were surely performing T2 was just before they provided an answer. They might have engaged in T2 at other times covertly, but we chose to study the impact of T2 on BLAST performance and T2-related neural processes in the windows preceding overt verbal responses.

One remaining issue is to estimate the duration of such windows: since the cognitive processes leading to the verbal responses are covert, there is no way to assess their duration and to specify a window during which participants are surely engaged in response selection for the VFT. Yet, a reasonable estimate could be derived from single-unit recordings performed during free-recall. Gelbard-Sagiv and colleagues observed that the activity of individual neurons in the human hippocampus increased about 1500 ms before the onset of the verbal report, when they recalled video clips they had seen recently and when a neuron with a strong response to that clip was being recorded (Gelbard-Sagiv et al, 2008). Since activity of individual neurons are thought to reflect the access to long-term memory in situations of recall, we considered the 2000 ms period preceding each verbal response as a wise choice to study T2-related processes (in the following, such windows will be called “T2-windows”).

Once T2 windows were defined, it became possible to evaluate if T2 interfered with T1 at the behavioral level, not only: a) by comparing of course the global performance (i.e. reaction time and accuracy) between the ST and DT conditions (referred to as “ST behavioral analysis”), but precisely, within the DT condition, b) by comparing performance during T2-windows and during windows with no verbal response (referred to as,“DT behavioral analysis”).

The “DT behavioral analysis” was performed as follows: in addition to the T2-windows (one window for every verbal response), all continuous time-segments free of any T2 responses and longer than 2s were divided into consecutive non-overlapping 2s windows called “T2-free windows”. Any difference in performance between T2 and T2-free windows was considered as a strong indication that T2 interfered with T1 at the behavioral level. Of course, participants might have engaged covertly in T2 during “T2-free windows”, but it is our best possible detection of T2-free periods and any misidentification would simply make our analysis too conservative. A straightforward randomization procedure was then designed to statistically estimate the impact of T2 on performance: i) first, the average reaction time (RT+) and accuracy (i.e. % or correct BLAST responses, AC+) were calculated across all T2-windows (N windows for N VFT verbal responses); B) the same procedure was performed on a random selection of N T2-free windows (excluding any window within 2s of a response) to compute a mean reaction time and accuracy value (RT-and AC-) for that selection; C) that last procedure was repeated 10,000 times to generate a distribution of 10,000 surrogate RT-and AC-values, to which RT+ and AC+ were compared, to provide two p-values, indicating of whether T2 had a significant negative impact on T1’s reaction time and accuracy (with a threshold of p=0.01).

### Electrophysiological Analysis

High-Frequency Activity between 50 Hz and 150 Hz (HFA[50-150] hereinafter, also termed ‘high-gamma’ by some authors) was extracted from iEEG time-series following our usual procedure. That procedure converts raw signals into time-series of neural activity which approximates neural spiking activity at the population level (Jerbi et al, 2009) and correlates tightly with behavior, even in real-time (Hamamé et al, 2012) and at the single-trials level (Ossandón et al, 2012).

#### Bipolar HFA Extraction

The data were recorded with a standard 128-channels acquisition system (Micromed, Treviso, Italy), bandpass filtered online from 0.1 to 200 Hz and sampled at 512 Hz in all patients. At the time of acquisition, the data were recorded using a reference electrode located in white matter, and the signal in each recording site was subsequently re-referenced to an adjacent channel on the same electrode (bipolar montage). A bipolar montage reduces signal artifacts common to adjacent electrode contacts (line-noise and distant physiological artifacts) and improves the spatial resolution of the recording to a few millimeters (Lachaux et al, 2003; Jerbi et al, 2009) — slightly from different from subdural grid electrocorticography (Jerbi et al, 2009) — by cancelling out effects of distant sources that spread equally to both adjacent sites through volume conduction. It might complicate functional connectivity analysis based on phase estimation,(Arnulfo et al, 2015) but no such analysis was performed here.

The frequency band of interest [50–150 Hz] was defined from preliminary time–frequency (TF) analyses of the iEEG data using wavelets (Tallon-Baudry et al, 1997), performed with an in-house software package for electrophysiological signal analyses (ELAN) developed at INSERM U1028, Lyon, France (Aguera et al, 2011), and from previous studies by our group (Jerbi et al, 2009). Raw data was extracted by using a homemade Matlab scripts (the Mathworks, Inc.) and transformed into HFA time-series with the following procedure (Ossandón et al, 2012; Perrone-Bertolotti et al, 2012).

Continuous iEEG signals were first bandpass-filtered in multiple successive 10 Hz wide frequency bands (e.g. 10 bands from [50–60 Hz] to [140–150 Hz]) using a zero phase shift no causal finite impulse filter with 0.5 Hz roll-off. The envelope of each bandpass-filtered signal was then computed with a standard Hilbert transform (Le Van Quyen et al, 2001), then down-sampled 64 Hz and divided by its means across the entire recording session and multiplied by 100, to express each value as a percentage (%) of that mean (normalization). Finally, the normalized envelope signals for each of the ten frequency band were averaged together to provide a single HFA time-series. By construction, the mean value of that HFA time-series across the entire recording session is equal to 100. The whole procedure is also designed to reduce the 1/f dropoff in amplitude of the raw iEEG signals. To detect task-related HFA increase or decrease induced by BLAST, the HFA time-series was epoched into data segments centered around BLAST target stimuli.

#### HFA Binarization

An additional analysis of HFA time-series was specifically developed to discriminate between “silent” vs “active” time-windows for any given recording site and to identify possible gaps left by T1 during which neurons could engage into T2. While a single-neuron can be said to be silent when it fires no action potential, the concept is difficult to transpose at the population level. In the specific context of this study, the best approximation for such “quiescent” state was during the baseline period of T1 (immediately before the onset of the Target). However, since T2 could potentially generate some intrinsic neural activity — this is our working hypothesis — we had to select episodes of relatively “pure” T1 activity.

For this reason, and considering the impact of T2 on behavioral performance — anticipating the results of the behavioral analysis provided in the results section — the baseline was chosen from the best 50% trials of T1 (the 50% fastest correct trials). Our best estimate of the “neutral” or “quiescent” state of a given neural population recorded with iEEG, was therefore the activity recorded in the best 50% T1 trials during a period (200 ms) immediately preceding the Target onset (shown in the white frame of Fig. 2A). The median of all HFA values measured during those windows — called Vmed — was computed for every iEEG site, and used to binarize the HFA signal: assigning a state ‘on’ or ‘off’ to every time-sample of the experiment. Fig. 2A and B illustrate that simple procedure, which was performed independently for each iEEG site. The end-result is a binary raster-plot, which reveals the impact of T1 and T2 on the local neural activity in the ST and DT conditions (Fig. 2B and D). Several statistical procedures were then applied to identify periods with enhanced activity due to T1 or to T2, and to study the interaction between the two.

**FIGURE 2.**
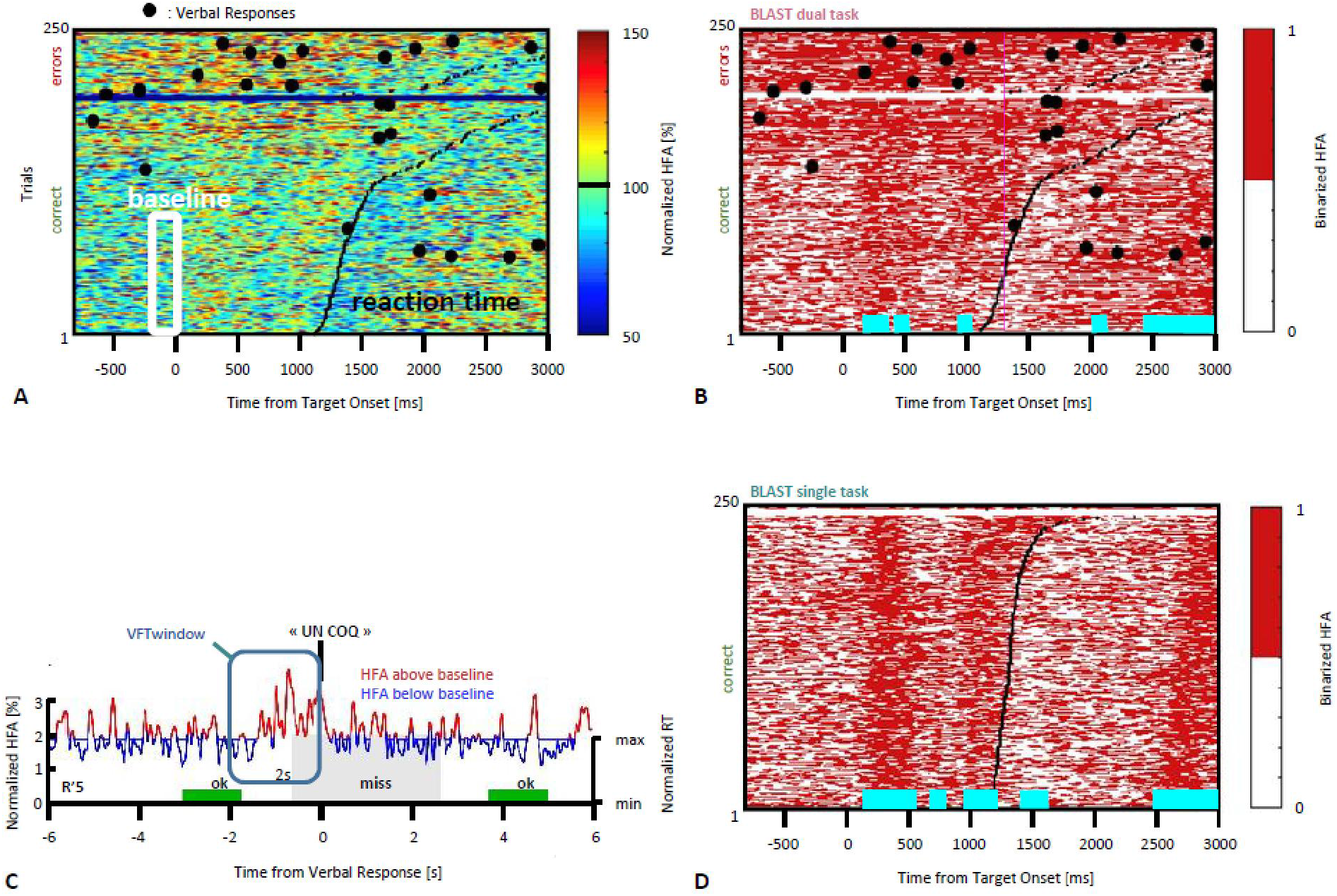
Methodological procedure. **A)** An example of neural response recorded in the left Precentral Gyrus during T1 in the DT condition, across all trials. Every row of the matrix corresponds to one trial, and HFA is color-coded as a function of time (x-axis). Trials are sorted by reaction time (black line) for correct and incorrect responses (in the lower and upper part of the matrix, respectively). HFA is expressed as a percentage of the average HFA value for that particular iEEG site, across the entire recording. Large black dots indicate verbal responses to T2. The white frame indicates HFA values used to estimate a quiescent baseline state (the prestimulus baseline during the 50 fastest correct trials of T1). The median of all HFA values within that white frame, Vmed, was used to create the binarized version of the matrix shown in B). **B)** binarized version of the same matrix (DT condition). By definition, ‘off’ samples in white (resp. ‘on’ samples, in red) have an HFA value below (resp. above) Vmed. During the baseline, the signal goes through a noisy succession of ‘on’ and ‘off’ states in equal number (at least, and by definition, for the best 50% trials); at other latencies, horizontal cyan lines indicate latencies with an abnormally high (or low) proportion of ‘on’ samples across the 50% fastest correct trials (see methods for details). By construction, cyan lines indicate time-windows during which the neural population is not available for T2. **C)** An example of HFA signal measured during a verbal response (“un coq” or “a rooster”). Values above Vmed (‘on’ samples) are shown in red, ‘off’ samples are shown in blue. Several of our analyses measured the proportion of ‘on’ samples within ‘T2-windows’, defined as the 2 seconds intervals before the response (vertical frame, in blue). T1 performance is color-coded for three consecutive trials (two hits in green, one miss in gray), and the height of the colored rectangles is proportional to reaction time (‘min’ = minimum reaction time across the entire experiment, same for ‘max’). **D)** Binarized trial matrix for the same iEEG site as in B) but in the ST condition. The comparison between B) and D) is illustrative of a strong interference between T1 and T2.

#### Statistical Analysis

##### Identification of T1-related neural responses

T1-related activity was detected from the binary raster-plots shown in Fig. 2B and D. When considering the 50% fastest correct trials for a given time sample in T1, (i.e. a vertical slice of the matrix, for instance the M values measured at the Target onset, at 200 ms, across the M fastest correct trials) an effect of the T1 task on neural activity in a given site should result in an abnormal number of ‘on’ samples relative to the prestimulus baseline. Such abnormalities were detected using a Wilcoxon test comparing that distribution of 0’s and 1’s with the distribution obtained for the [-200 ms : 0 ms] baseline (in all figures, time-segments with a significant deviation from the baseline are indicated by a horizontal cyan lines below the plots; with a statistical threshold of p<0.05 and a False-Discovery Rate correction for multiple comparisons).

##### Identification of T2-related neural responses

Binarized HFA signals were also used to identify iEEG sites in which T2 had an effect on the local neural activity. Using the same logic as for the behavioral analysis, a T2-window of 2s was defined for each of the N verbal response of the participants. Then, the percentage of ‘on’ samples was calculated across N T2-windows (PCT+), and for N randomly chosen T2-free windows (PCT-), repeating the surrogate procedure 10,000 times to create a distribution of 10,000 surrogate PCT-values, to which PCT+ was compared to obtain a p-value (and apply a statistical threshold of p<0.05, Bonferroni-corrected for multiple comparisons, as the FDR correction could not be applied here). In the Results section, iEEG sites identified with that procedure are said to have an “abnormally dense activity” during T2.

##### Detection of sites of interference between T2 and T1

To identify iEEG sites in which T2 interfered with T1, the analysis was based on the observation that T1 trials repeat roughly every 2 seconds: 800 ms of fixation followed by 200 ms of Target presentation, a 500 ms mask and roughly 500 ms before the earliest button presses (Fig. 1B). Given that convenient value, similar to the duration of T2-windows, an estimate of the typical density of ‘on’ states during optimal T1 performance could be estimated from the fastest correct trials, by calculating the proportion of ‘on’ sites in a [-800 ms : 1200 ms] surrounding the Target onset. Any evidence that the density of ‘on’ state during T2-windows was greater than during “high performance” T1 windows was indicative of a possible interference between the two tasks at the neural level, with an excess of neural activity due to T2. The actual comparison used a Wilcoxon sign-rank test to compare between the density of ‘on’ states measured in the N T2-windows (N density values ranging between 0 and 1, i.e. one density value for each T2-window) and the density of ‘on’ states measured in the N T1 windows corresponding to the N fastest correct trials. The result was an “Interference detection test” which identified all iEEG sites with a higher density of ‘on’ states during T2-windows (p<0.05, Bonferroni correction).

##### Comparison of T1-related activity between the ST and DT conditions

The direct comparison of HFA signals recorded in separate sessions can be biased by line-noise variations, even if a bipolar montage reduces such contamination. By comparing the binarized HFA times series recorded in the ST and DT sessions, we reduced that bias: our analysis compared the proportion of ‘on’ values in the M best trials of the ST and DT conditions for every sample within a [-1000 : 3000 ms] interval surrounding target onset (T = 0 ms), where M was set to half the number of correct trials in the DT session (i.e. considering the 50% fastest correct trials of BLAST to minimize the effect of T2). A Wilcoxon test was used to compare for each sample the distributions of M activity values (0’s or 1’s) for the ST and DT conditions (p<0.05, with a False-Discovery Rate correction for multiple-comparisons).

Finally, a similar procedure was used to test whether the activity of specific regions involved in executive control varied between the beginning and the end of the ST session of BLAST. The motivation was to test for a possible automatization of BLAST throughout that initial session (see results). The analysis was as previously described, except that the comparison was performed between the first and last 20% trials (50 trials for each group).

### Correlation Analysis

For every pair of iEEG sites, and for every VFT verbal response, we considered a 10s window centered on the response [-6000 : 4000 ms; response = 0 ms] and calculated the correlation coefficient (R+) between the two HFA time-series for that window. R+ was compared to a population of 10,000 surrogate correlation coefficients created by repeating the following procedure 10,000 times: a) consider two random iEEG sites separated by more than 30 mm and for each of them, a 10s window around a verbal response (with no temporal overlap between the two windows), then b) compute the correlation coefficient R-between the corresponding HFA signal, c) repeat for i = 1 to 10,000. The initial R+ coefficient was considered to be significant if greater than all surrogate R-values (p < 0.0001, corresponding to a conservative Bonferroni correction). This procedure was used to identify VFT responses with a significant correlation between two iEEG sites.

### Anatomical Display

The precise anatomical location of the electrodes (and their MNI coordinates) were obtained by aligning each patient’s pre and post-implantation MRIs using the NUTMEG toolbox (Dalal et al, 2004) and IntrAnat (Deman et al, 2018), a specific toolbox interfacing with the BrainVisa software (IntrAnat Electrodes, GIN INSERM U1216, Grenoble, available at https://f-tract.eu/index.php/softwares/). Participants’ behavior was recorded and inspected using an in-house software — BrainTV Replay — while anatomical representations were made using HiBoP, also developed by our group and available here (https://github.com/hbp-HiBoP/HiBoP/releases/tag/2.2.3a).

BrainTV Replay is an updated version of our original BrainTV software.(Lachaux et al, 2007, 2012; Vidal et al, 2011) It is designed to visualize a “replay” of the entire experimental session for any participant, with a simultaneous display of the video, the behavioral responses and HFA activity measured for all iEEG sites (Fig. 1A and Fig. 11) both as time-series and onto a 3D representation of the patient’s brain. BrainTV Replay also allows the rapid visual identification of highly correlated HFA fluctuations between iEEG sites. HiBoP was designed to visualize iEEG signals onto 3D anatomy at both group and single-patient level (Fig. 1E). It allows to label iEEG sites using cortical parcellations such as Brodmann’s or Marsatlas (Auzias et al, 2016) and the Freesurfer/BrainVisa pipeline (available at http://surfer.nmr.mgh.harvard.edu) (Fischl, 2012) (see Supplement material: Table I)

## RESULTS

### Behavioral Analysis

Our main assumption was that high performance in T1 (i.e. fast reaction times, few errors) requires a specific dynamic pattern of cortical activations and that T2 might interfere with that pattern in the DT condition to decrease performance. Accordingly, the T1 response selection process should be delayed (slower reaction time) or cancelled (the participant would either not respond or take a bet with a 50% chance of failure) when performing T2 concurrently.

The results of the “ST behavioral analysis” (Fig. 1D) largely confirmed that prediction at the global performance level: overall, reaction times were slower and accuracy lower in the DT than in the ST condition (with a significant increase in reaction time in 11 of the 12 participants). The more detailed “DT behavioral analysis” also revealed that T1 performance in the DT condition was impaired when participants actively searched for T2 answers (all patients had either a significantly slower reaction time or a lower accuracy in periods immediately preceding verbal responses, as exemplified in Fig. 2C). Overall, the behavioral data obtained in the ST and DT conditions largely confirmed the difficulty for patients to perform both tasks at the same time, which motivated the subsequent search for interferences between the two tasks at the neural level.

### Electrophysiological Analysis

Marked differences were observed between T1 activation patterns in the ST and DT conditions. As an illustrative example and an introduction to our analysis strategy, Fig. 1C shows the response in the left Precentral Gyrus to all T1 trials in the two conditions, sorted by reaction time and accuracy. Besides the higher error rate in the DT situation, the most visible difference is the presence of sustained blocks of activation in DT, which are largely absent in ST and incompatible with the fine temporal structure of the neural activation pattern supporting T2 (as visible for the fastest correct trials).

To identify cortical sites with a similar effect, we ran the interference detection procedure described earlier to detect iEEG sites with increased HFA before T2 verbal responses (“T2-windows”), as compared to other time periods of the DT session, including the best T1 trials. We found that the optimal T1 electrophysiological response was disrupted in DT in six major cortical areas: the left Dorso-Lateral Prefrontal Cortex (DLPFC); the Lateral Temporal Cortex (LTC); the Anterior Insula bilaterally (AI), the junction of the Anterior Cingulate Cortex and the Pre-Supplementary Motor Area (ACC/preSMA), the Precentral Gyrus/Sulcus (Precentral cluster), and the Basal Temporal Cortex (BTC). We now discuss each cluster separately.

#### Precentral Cluster

We found effects in the Precentral Gyrus (1 LH, 1 RH, 2 patients) and in the Precentral Sulcus (2LH, 1 RH, 3 patients) with similar neural responses in all sites. The optimal T1 response was characterized by two separate phases of activation until the manual response (Fig. 3). In the DT condition, a strong and sustained activation was mostly present during the bad trials (i.e. slowest reaction time and/or higher error rate). Recordings from the same sites during additional tasks (the functional localizers) revealed a strong activation during a rhyming task emphasizing phonological processing, and during a verbal working memory task (data not shown) (Vidal et al, 2011). These are strong indications that our electrodes sampled a region specialized in covert speech and verbal rehearsal. That might explain the local interference between T1 and T2, because the participants seem to have used a verbal strategy to encode the T1 Target, while the search for specific words in the latter also requires a covert linguistic production. The observed disturbance could most likely be interpreted as direct evidence of a specialized bottleneck, involving a specific cognitive component shared by the two tasks. Yet, we also found clear evidence for a strategy change during T1 for some patients, who seem to have used a non-verbal strategy during the best trials, with a strong attenuation of the double activation peak (Fig. 3).

**FIGURE 3.**
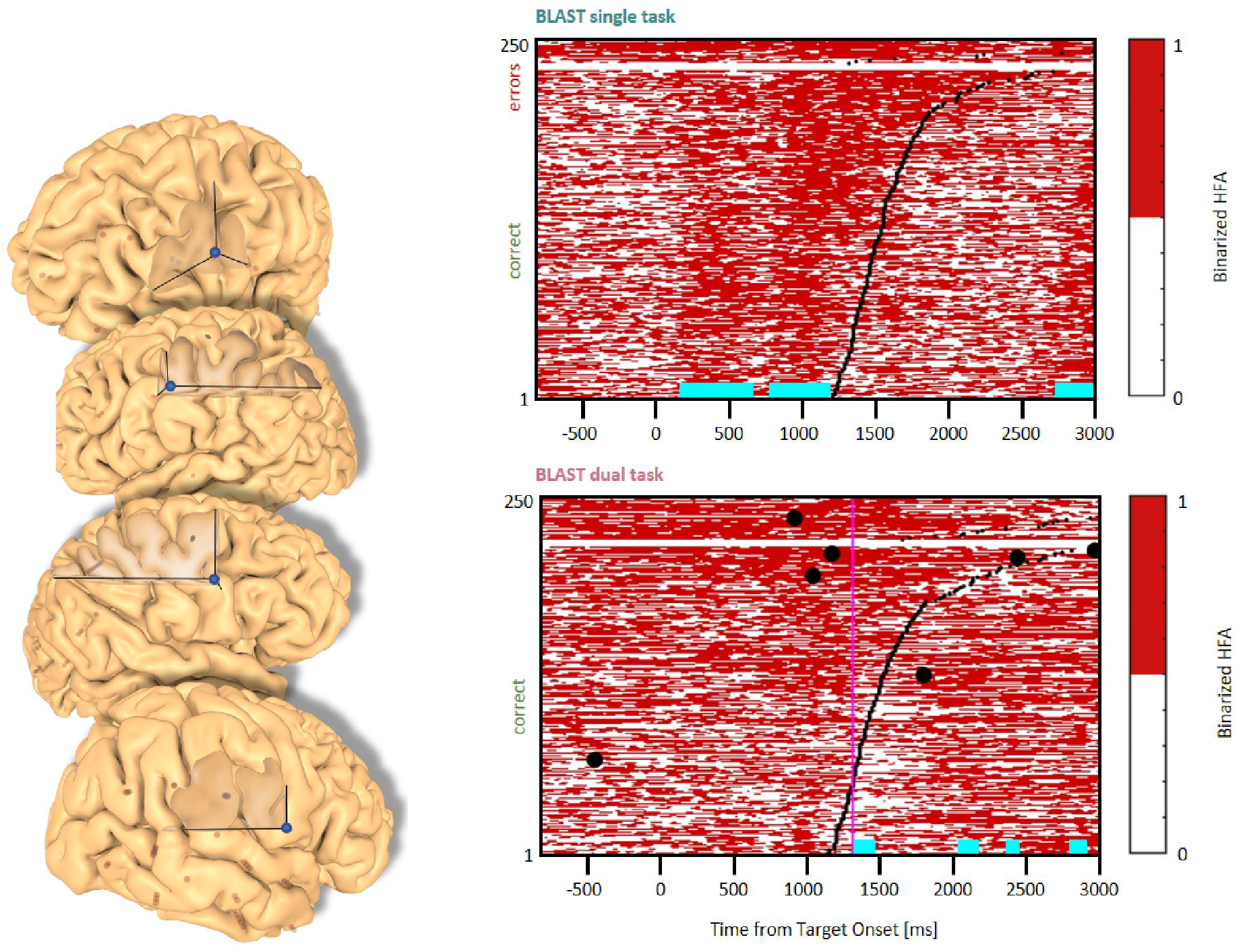
Interference in the Precentral Cluster. Sites with an interference between T1 and T2 are displayed at their precise anatomical location onto a 3D representation of the participant’s brain. The matrix representation reproduces the display of Fig. 2, for the single-task and dual-task conditions for one of the sites. Patients id [Px] and iEEG site names, from left to right and top to bottom: R’9 [P4]; S’9 [P5] (**matrix displayed**); M9 [P2]; R9 [P1].

#### DLPFC Clusters

Two types of effects could be distinguished in the lateral prefrontal cortex. In the Inferior Frontal Sulcus immediately adjacent to Broca’s pars triangularis, a double activation peak was observed in three sites (LH, 3 patients) while participants encoded the Target and searched the Array (Fig. 4), very much as it was observed in the Precentral cluster. In all three sites, the interference in the DT condition produced a response attenuation for the fastest correct trials, and a contamination by T2 during the slower and/or incorrect trials, characterized by a sustained activation. In separate verbal localizers, that sub-cluster was particularly active during semantic and phonological processing, as well as during the retrieval of items stored in verbal working memory. This is consistent with a role in manipulating verbal working memory, information as required by T2 and T1 when the strategy is verbal. The response attenuation in the best trials of the DT condition would again indicate to a shift towards a different strategy, in which the Target would be encoded as a visual template — possibly — rather than in a verbal, phonological form (Rogers & Monsell, 1995). An interference pattern was also observed at three other sites more anterior in the DLPFC, specifically in the Middle Frontal Gyrus above the anterior pars triangularis (LH, 3 patients, Fig. 5). There, T1-related activity was different from the previous pattern, with no response during the Target presentation and an activation by the end of the Array presentation extending to the short inter-trial pause. The role of that sub-region in both tasks is less easy to understand, but the corresponding sites were selectively activated by language and verbal working memory localizers.

**FIGURE 4.**
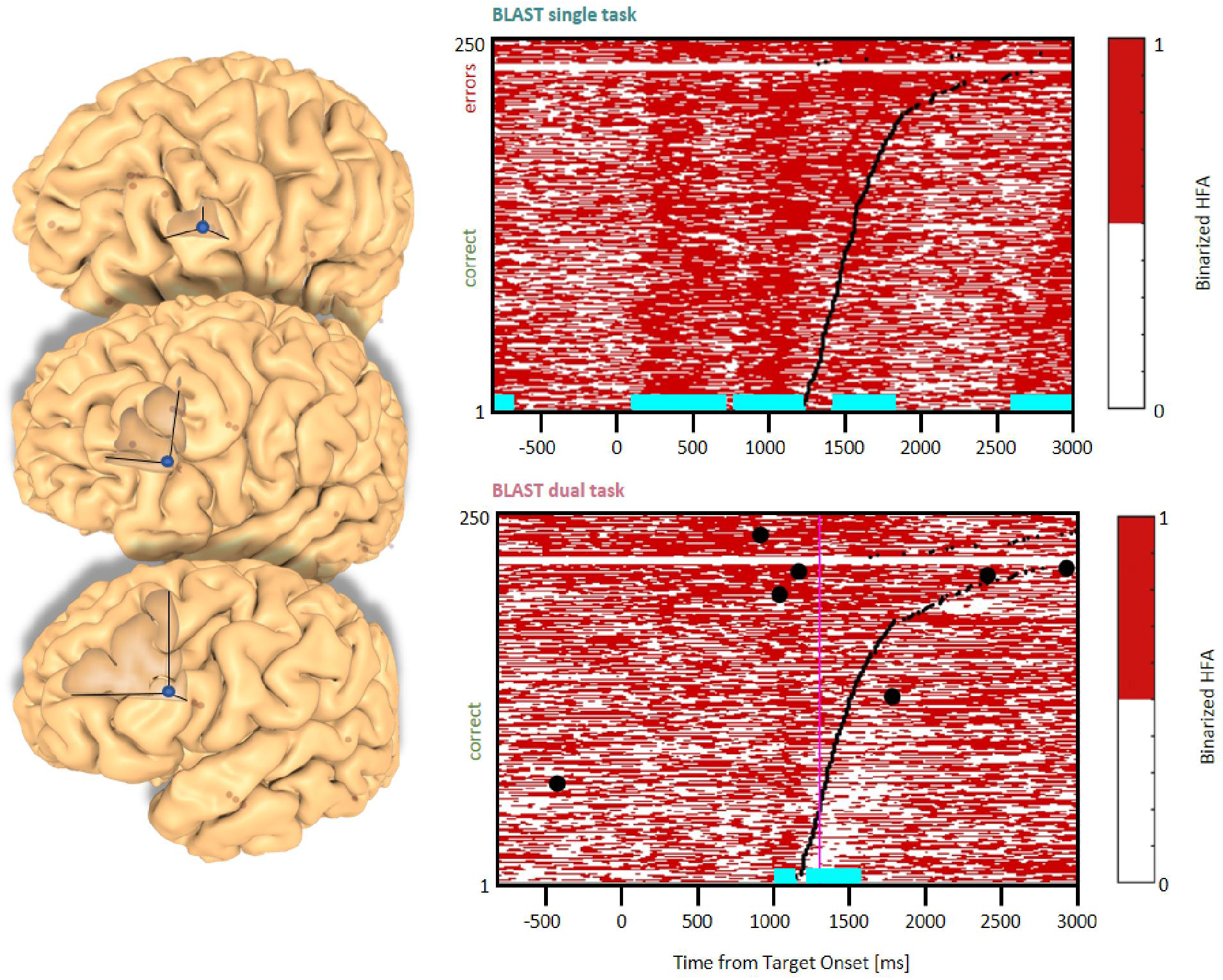
Interference in the Inferior Frontal Gyrus/Sulcus. Sites with an interference between T1 and T2 are displayed at their precise anatomical location onto a 3D representation of the participant’s brain. The matrix representation reproduces the display of Fig. 2, for the single-task and dual-task conditions for one of the sites. Patients id [Px] and iEEG site names, from left to right and top to bottom: G’13 [P4]; K’10 [P5] (matrix displayed); Q’2 [P11].

**FIGURE 5.**
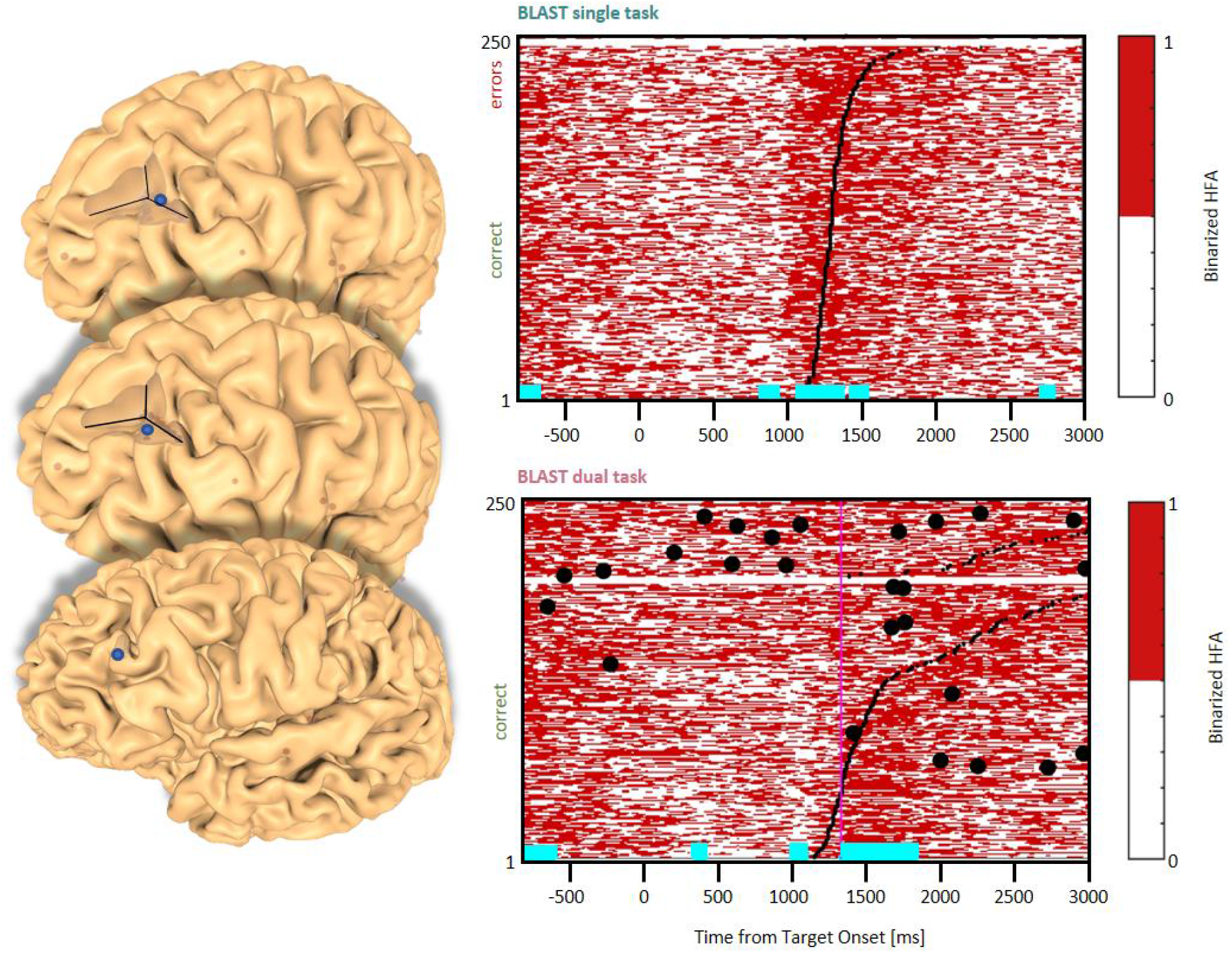
Interference in the Dorsolateral Prefrontal Cortex. Sites with an interference between T1 and T2 are displayed at their precise anatomical location onto a 3D representation of the participant’s brain. The matrix representation reproduces the display of Fig. 2, for the single-task and dual-task conditions for one of the sites. Patients id [Px] and iEEG site names, from left to right and top to bottom: F’9 [P4] (matrix displayed); Y’13 [P4]; G’12 [P9].

#### Anterior Insula & ACC/preSMA Clusters

In the Anterior Insula, we found six sites (2 in the LH, 2 patients, and 4 in the RH, 4 patients) with a deviation from the optimal T1 dynamics in the DT condition (Fig. 6). The T1 response pattern was reproducible across sites, with a sustained activation during the display of the Array, time-locked to the response. T2 also produced sustained activations, clearly incompatible with the T1 activation pattern. Both left and right hemisphere sites were activated by a verbal working memory task and by tasks emphasizing semantic and phonological processing. These observations were mirrored by responses in the Paracingulate Sulcus (one site in each hemisphere, 2 patients, Fig. 7), although the activity rose more progressively throughout T1 trials. Data from the functional localizers revealed a similar increase in all tasks, emphasizing executive control (visual search, working memory). As often seen before, the neural activation during the best T1 trials (i.e. 50% fastest correct trials) was weaker or even absent in the DT conditions, compared to the ST condition (Fig. 12). That specific result should catch our attention and will be discussed further, considering that this specific anatomical cluster is believed to support general executive control, rather than specific cognitive processes: it suggests the possibility of a general change of approach to cope with the DT situation, much more global than a simple strategy change from a verbal to a visual encoding of the Target, for instance.

**FIGURE 6.**
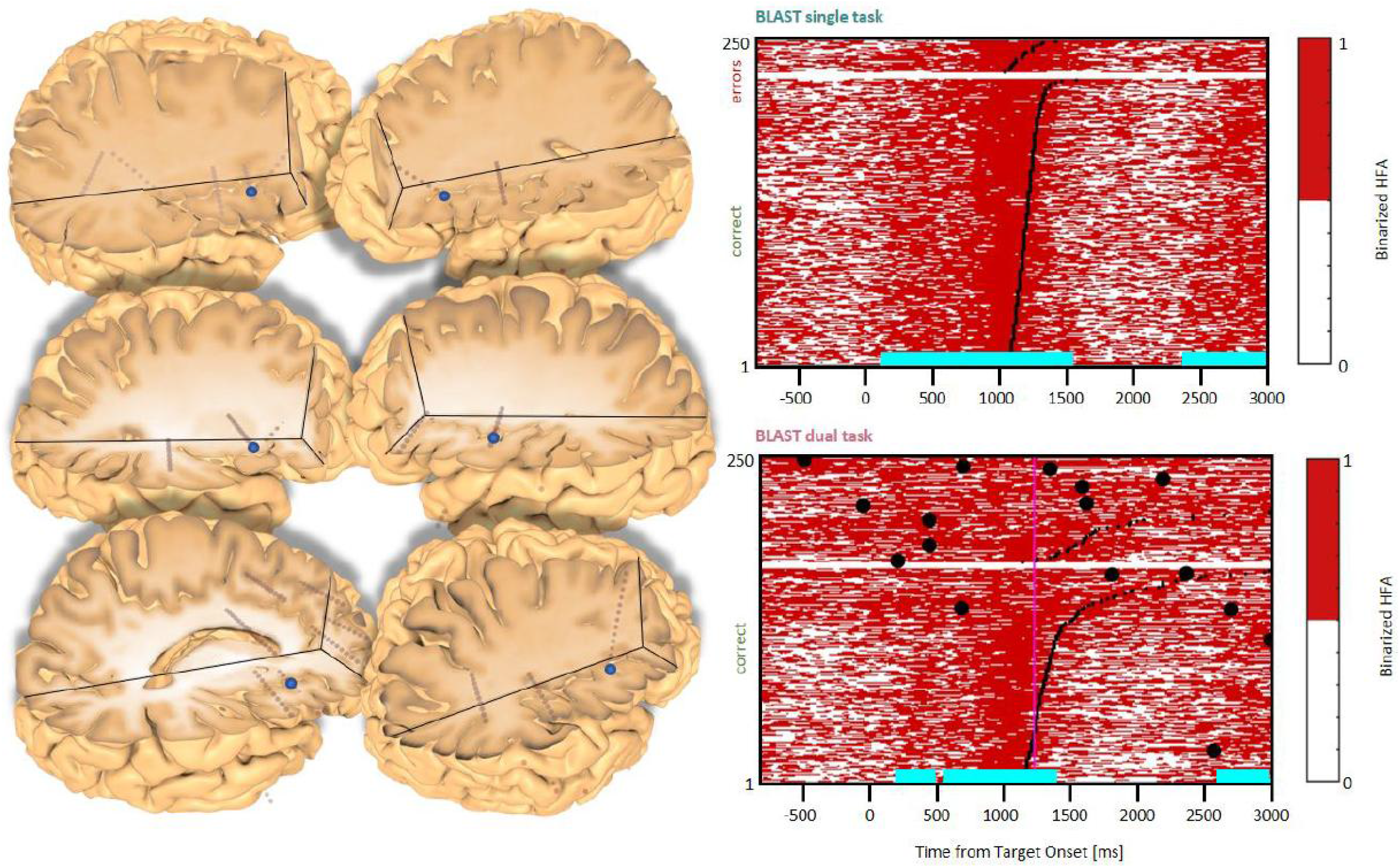
Interference in the Anterior Insula. Sites with an interference between T1 and T2 are displayed at their precise anatomical location onto a 3D representation of the participant’s brain. The matrix representation reproduces the display of Fig. 2, for the single-task and dual-task conditions for one of the sites. Patients id [Px] and iEEG site names, from left to right and top to bottom: X7 [P3]; X’9 [P3]; X5 [P11]; **X’9 [P11] (matrix displayed)**; E4 [P7]; X7 [P12].

**FIGURE 7.**
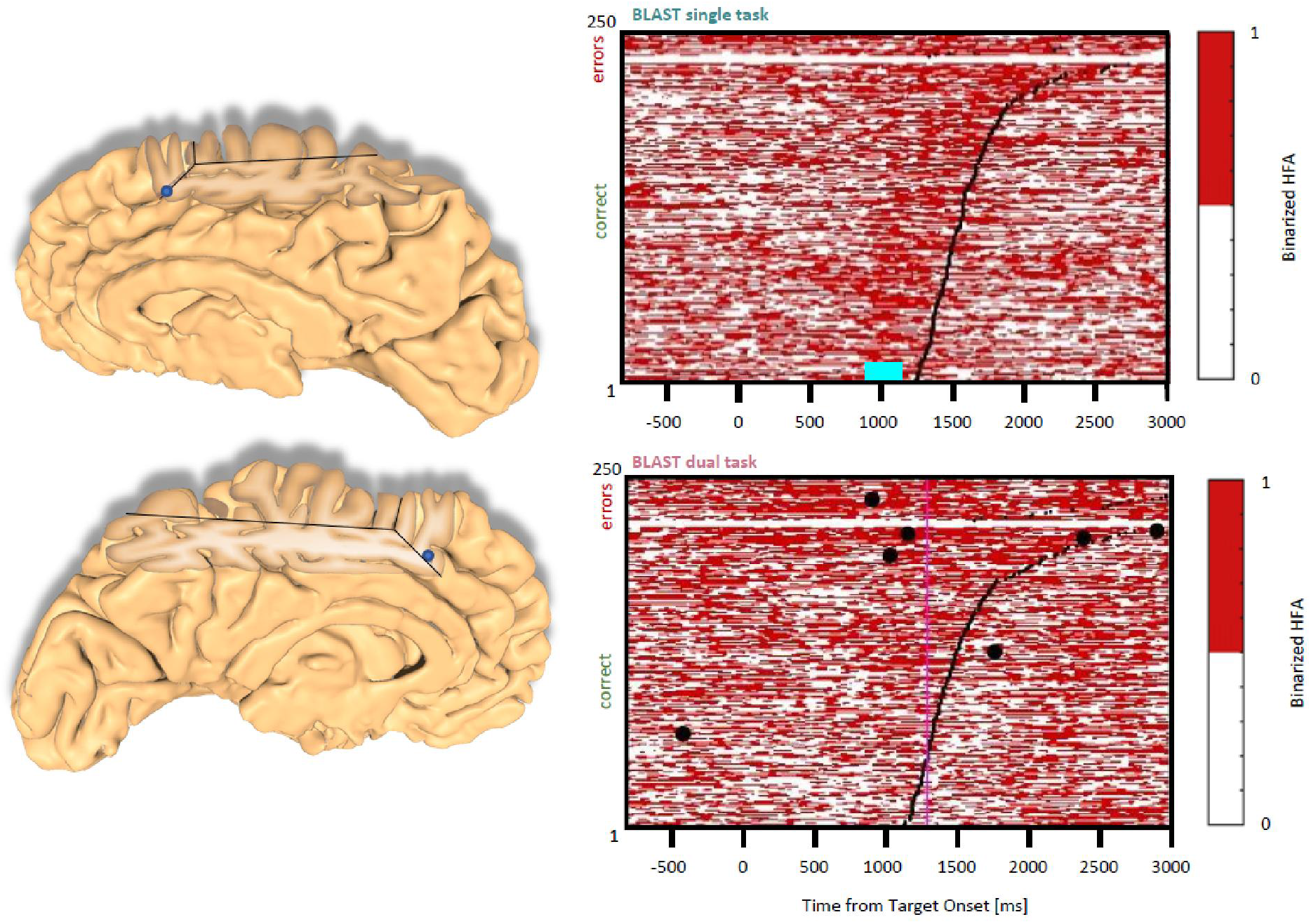
Interference in the Anterior Cingulate Cortex and the Pre-Supplementary Motor Area. Sites with an interference between T1 and T2 are displayed at their precise anatomical location onto a 3D representation of the participant’s brain. The matrix representation reproduces the display of Fig. 2, for the single-task and dual-task conditions for one of the sites. Patients id [Px] and iEEG site names, from left to right and top to bottom: S2 [P7]; **Z’2 [P5] (matrix displayed)**.

#### Lateral Temporal Cluster

Three sites activated by T1 in the Middle Temporal Gyrus were also reactive to T2 (Fig. 8, two sites in the left hemisphere and one in the right hemisphere in two patients). The activation patterns were reminiscent of the Precentral cluster, both for T1 and for the functional localizers, with strong and selective responses to conditions emphasizing verbal processing. We suggest that this cluster might support verbal processes shared by T2 and T1 under a specific verbal strategy. The sharp reduction in activity in the best T1 trials of the DT condition could again be explained by a shift to a non-verbal strategy to reduce interference. Once again, the amount of interference between T1 and T2 seems to depend more on the strategy used rather than on the tasks themselves.

**FIGURE 8.**
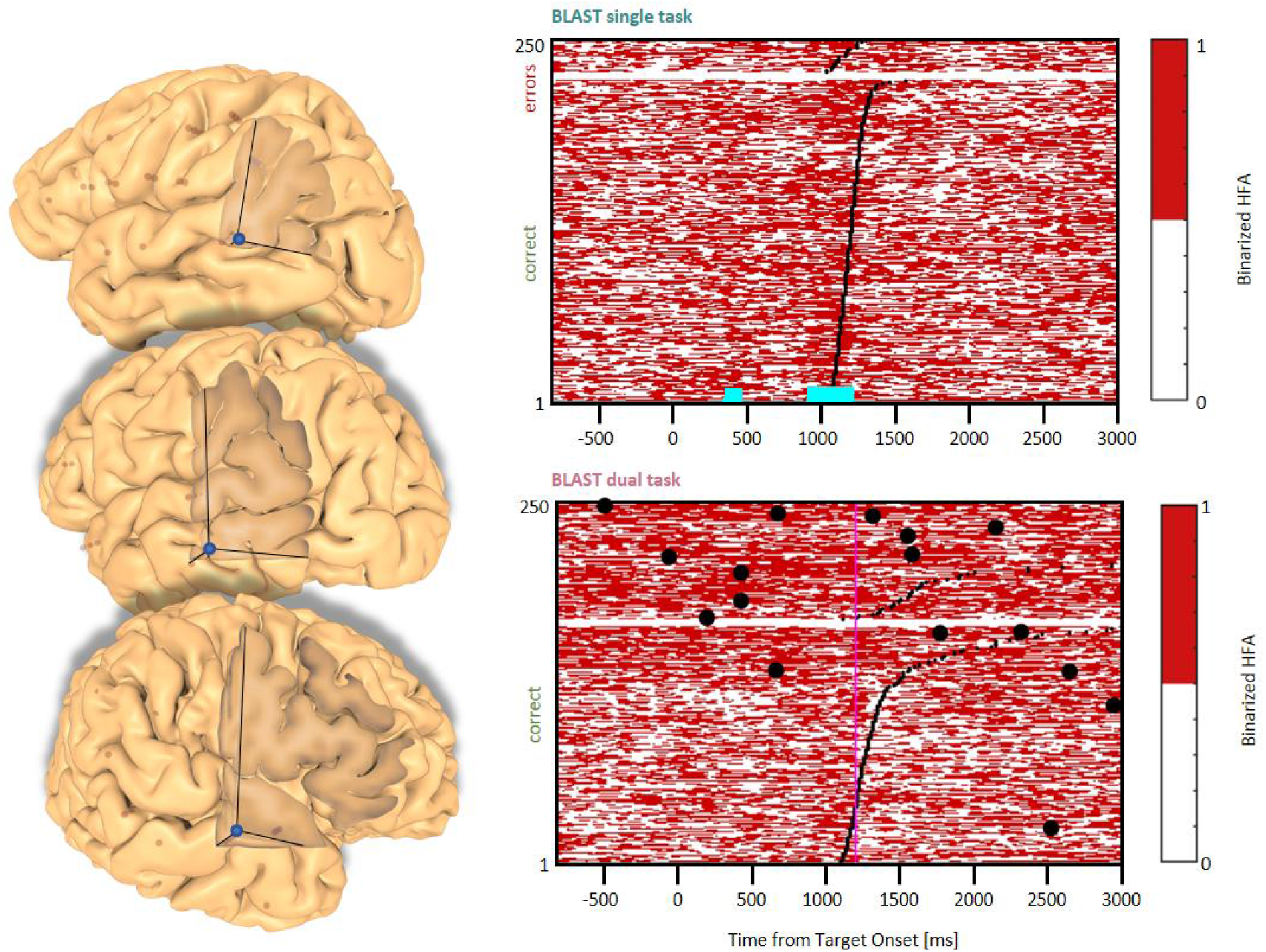
Interference in the Lateral Temporal Cortex. Sites with an interference between T1 and T2 are displayed at their precise anatomical location onto a 3D representation of the participant’s brain. The matrix representation reproduces the display of Fig. 2, for the single-task and dual-task conditions for one of the sites. Patients id [Px] and iEEG site names, from left to right and top to bottom: D’7 [P6]; **C’12 [P11] (matrix displayed)**; A9 [P12].

#### Basal Temporal Cluster

Finally, an interference pattern was also observed in three sites around the Fusiform Gyrus (2 in the LH, 1 in the RH) (Fig. 9). Interestingly, all three sites were located in sub regions of the visual cortex specific to precise object categories, as revealed by responses to a functional localizer showing several categories of pictures. Both sites in the left hemisphere were located in the Visual Word Form Area (selective to letter strings), while the right hemisphere site was most efficiently activated by face stimuli (but still responsive to letter strings). The activation of these sites during T2 was not unexpected, since participants could rely on mental imagery to remember specific faces, find names starting with the given letter, or visualize spelling. Still, it is quite remarkable that such activity could be detected online, revealing in real-time the strategy used by the participant. Incidentally, T2-related activities turned out to be stronger in the right hemisphere when the participant was searching for people’s names vs. other categories, such as fruits.

**FIGURE 9.**
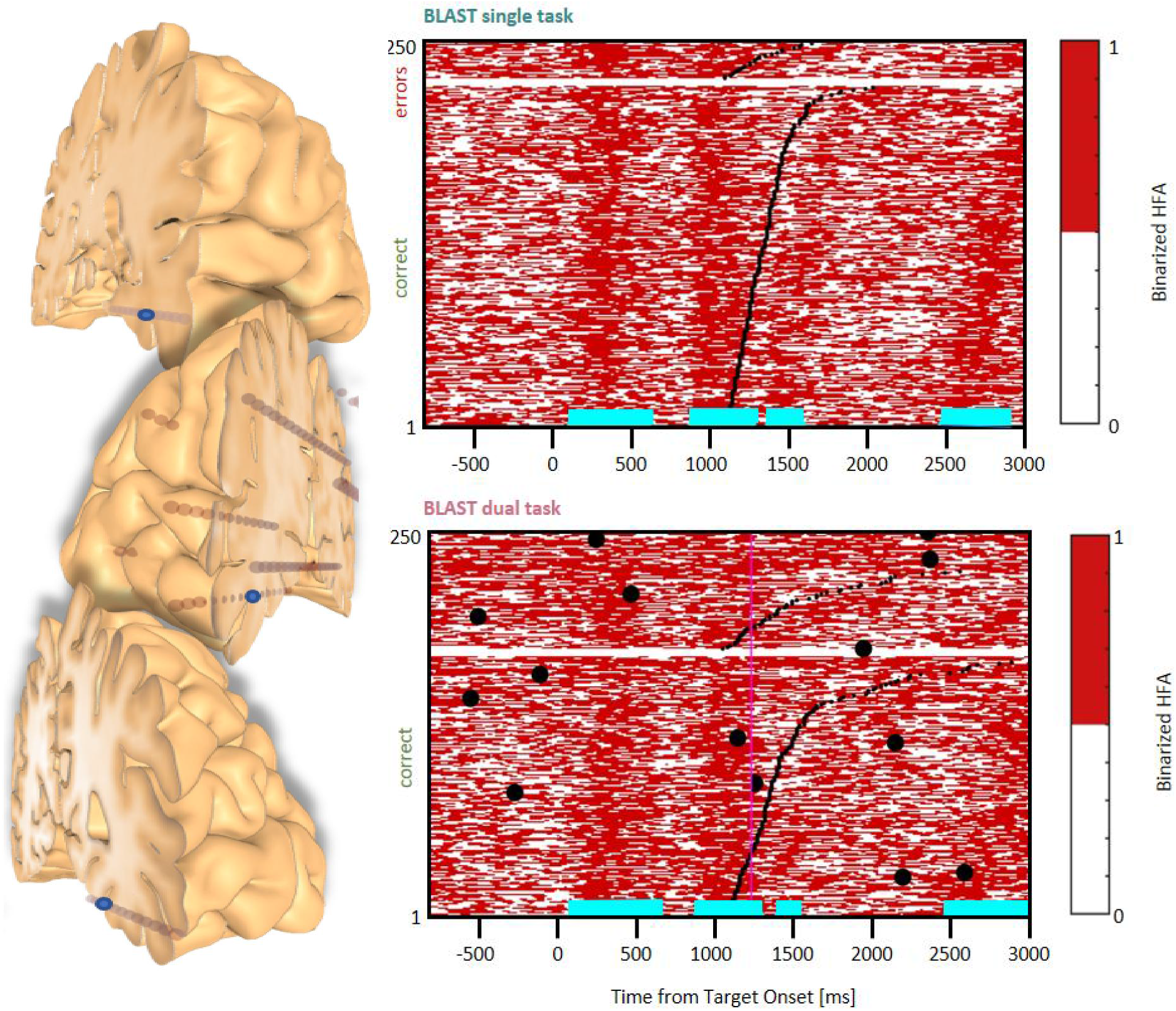
Interference in the Basal Temporal Cortex. Sites with an interference between T1 and T2 are displayed at their precise anatomical location onto a 3D representation of the participant’s brain. The matrix representation reproduces the display of Fig. 2, for the single-task and dual-task conditions for one of the sites. Patients id [Px] and iEEG site names, from left to right and top to bottom: E’7 [P4]; **F6 [P1] (matrix displayed)**; L’3 [P5].

### Response Selection is slowed down, not postponed

Considering the intense debate regarding whether a secondary task can postpone the response selection process of a primary task, we paid a specific attention to two patients with a clear BLAST response component visible in sites with no significant contamination by the VFT (Fig. 10, top four panels, P4 G’8 and P5 S’12, respectively in the middle frontal gyrus and the precentral gyrus). Because of the absence of direct VFT interference, the response dynamics was clear in both ST and DT conditions, and indicative of a response selection process (i.e. sustained from the stimulus to the response). Notably, the dominant effect in the DT condition was a longer, not later, activity in both sites, for the slower (‘bad’) trials. Accordingly, the response selection process appeared to be slowed down, rather than postponed. Corresponding neural correlates of the response selection process are shown for two VFT-contaminated sites (Fig.10, bottom four panels, P4 R’4 and P5 K’10, for comparison, respectively in the precentral sulcus and the inferior frontal sulcus).

**FIGURE 10.**
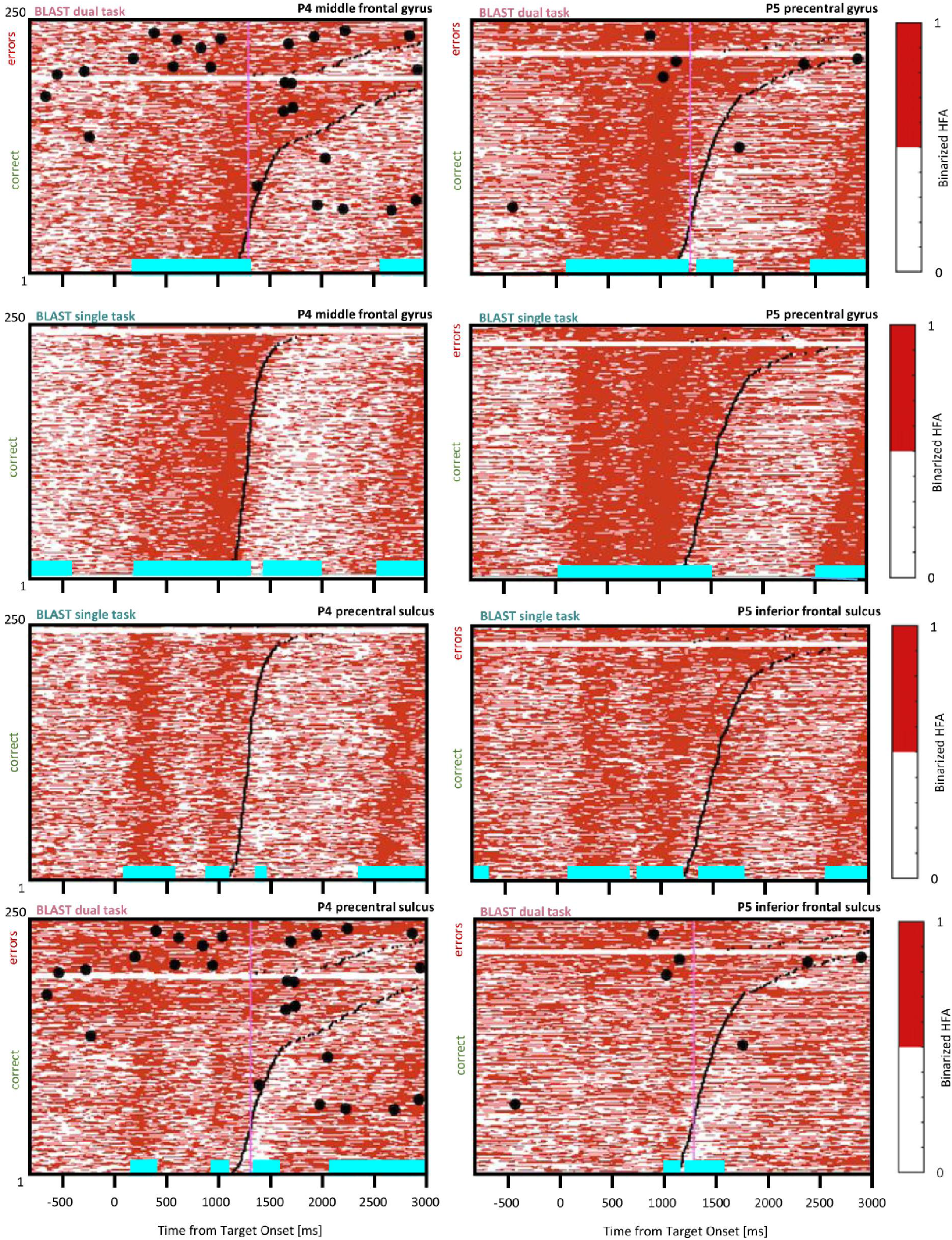
Slowing down of the response selection process. Comparison of BLAST-related activity in the ST and DT conditions for four sites (two of them with no significant VFT interference - top four panels). For each site, BLAST-related neural activity is represented as a binarized matrix for the ST and DT conditions. Horizontal lines (cyan) indicate samples with a significant increase or decrease relative to the [-200:0 ms] pre-Target baseline. From left to right and top to bottom, iEEG sites and [Patients id] are : G’8 [P4]; S’12 [P5]; R’4 [P4]; K’10 [P5].

### Network Analysis (and the possibility of Indirect Interference)

Our visualization software, BrainTV Replay, allows for the joint review of HFA times series of two sites, as superimposed curves synchronized with video recordings, which is extremely convenient to detect high-correlation episodes visually (Fig. 11). During the preliminary visual review of the data, we were immediately struck by extremely high correlations between HFA time fluctuations of specific sites distant from each other. This motivated a quantitative search for significant functional coupling between the sites reported above, when several clusters could be recorded in the same patient. The analysis revealed patterns of very high and significative correlation episodes for specific verbal responses and cortical areas (as illustrated between the left Precentral cortex and the left BTC in Fig. 11, left panel); however, few of them were systematic enough to be reported. The most reproducible correlation was observed in two patients (P3 and P11), between the left and right Anterior Insula, where we found significant correlations between both hemispheres for more than half of T2’s verbal responses (13 out of 16 for P3, and 17 out of 31 for P11) (see Fig. 11 right panel for an example). Although limited, our results showed that “the” central bottleneck should rather be thought as a network of bottlenecks, where tasks interact dynamically with each other. This also raises the possibility that a secondary task like T2 could also interfere indirectly with T1, by recruiting cortical regions that do not belong to the T1 network, but that are coupled to some of its components and interact with T1 through that interaction.

**FIGURE 11.**
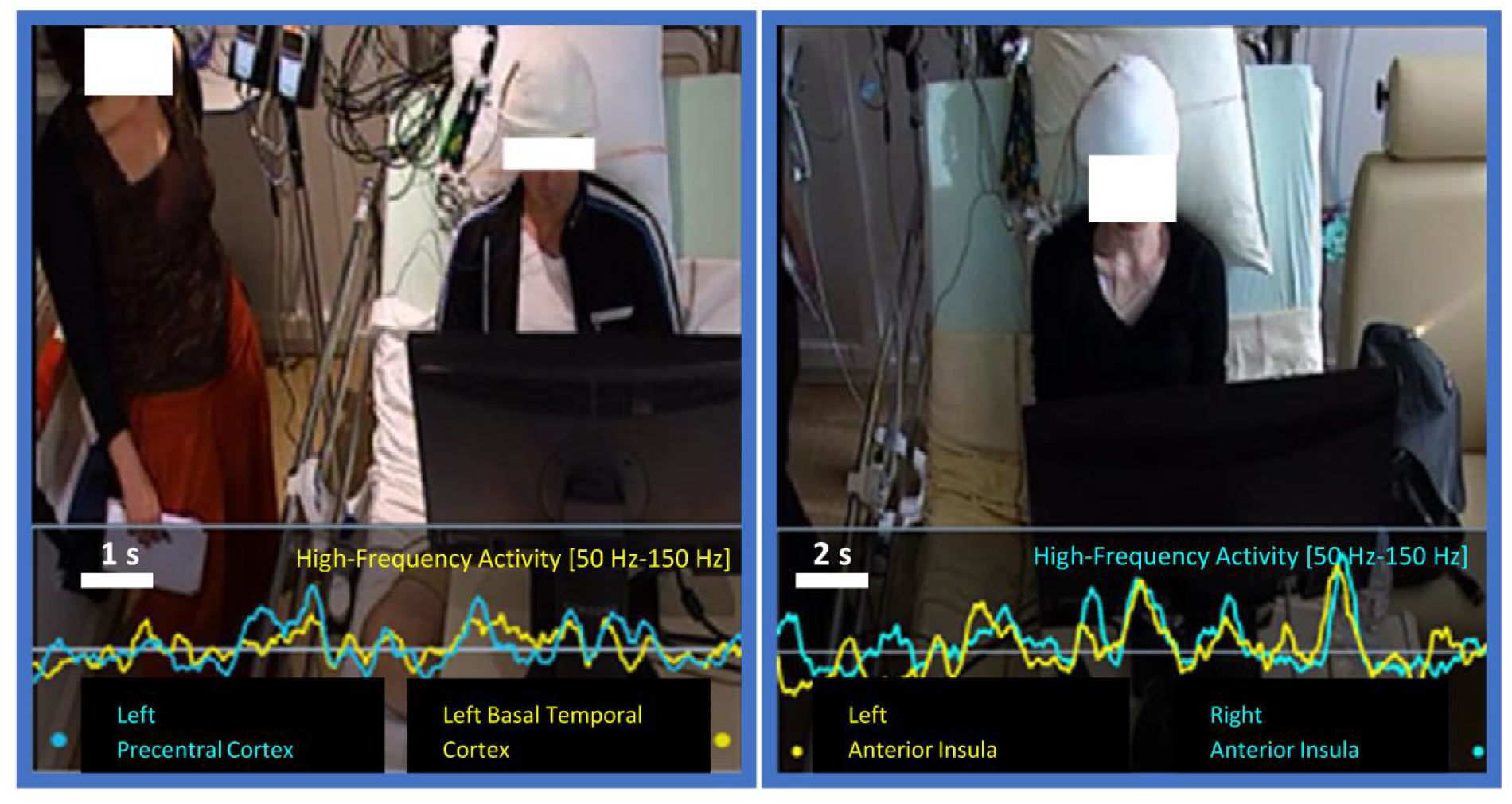
Correlation between bottlenecks. A) BrainTV display showing an episode of very high and sustained correlation between two bottlenecks, one in the Precentral Cluster and the other in the Basal Temporal Cluster. Curves correspond to HFA activity measured in the last 10 or 20 seconds before a verbal VFT response, as indicated in the right panel). Same representation showing a robust correlation between two sites in the left and right Anterior Insulae.

### Fine Tuning of Executive Control

To conclude, we specifically investigated the differential involvement of the executive control network in the ST and DT conditions. As reported above, some sites in the DLPFC, the AI and the ACC/preSMA were less active during the best trials of BLAST in the DT than in the ST condition, as recapitulated in Fig. 12. The figure reveals a significant difference between the density of ‘on’ states in the two conditions, when considering the same number of trials with the fastest correct responses (see methods). In some sites situated in essential nodes of a network supporting cognitive control, an attention-demanding task such as BLAST evoked almost no response in the DT condition, which comes as a surprise. We reasoned that participants might have automatized BLAST through the ST session, which always came before the DT session; in which case an attenuation of the response to BLAST should already be visible within the ST session. To test that hypothesis, we tested whether activity in the executive network was lower at the end than at the beginning of the ST session. For all sites discussed in this section, we measured the density of ‘on’ states for each trial, in a critical [900 : 1200 ms] window relative to the Target onset, and compared the values obtained for the first 20% of the trials (50 trials at the beginning of the ST session) and the last 20%. We found no significant difference except for one site in the ACC where activity was greater by the end of the experiment. It is therefore unlikely that the reduced response of the executive control network in the DT condition was caused by the automatization of BLAST. It is most likely due to an effective reduction of control with no performance decrement, which suggests that participants exerted more control in the ST condition than strictly necessary.

**FIGURE 12.**
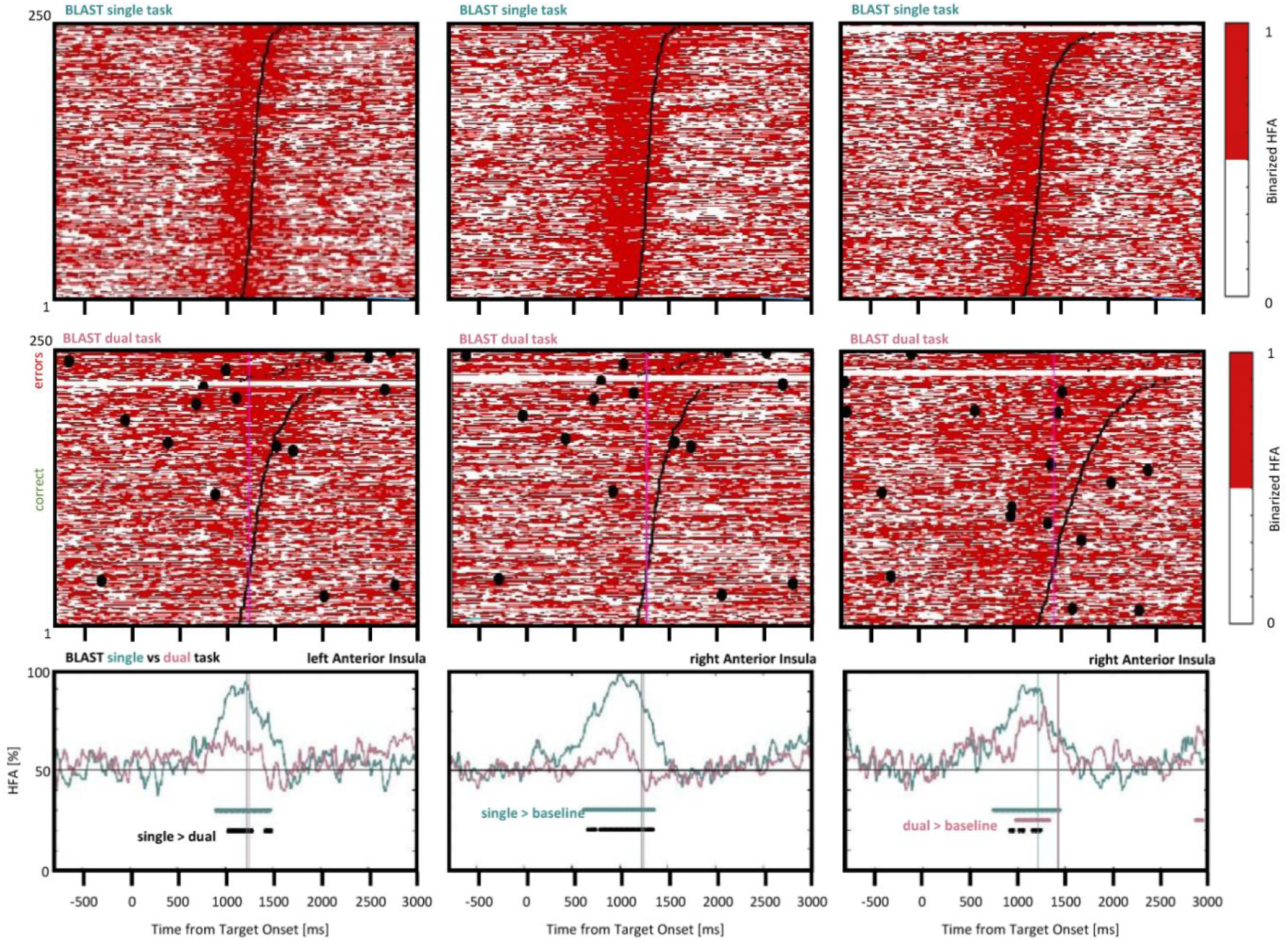
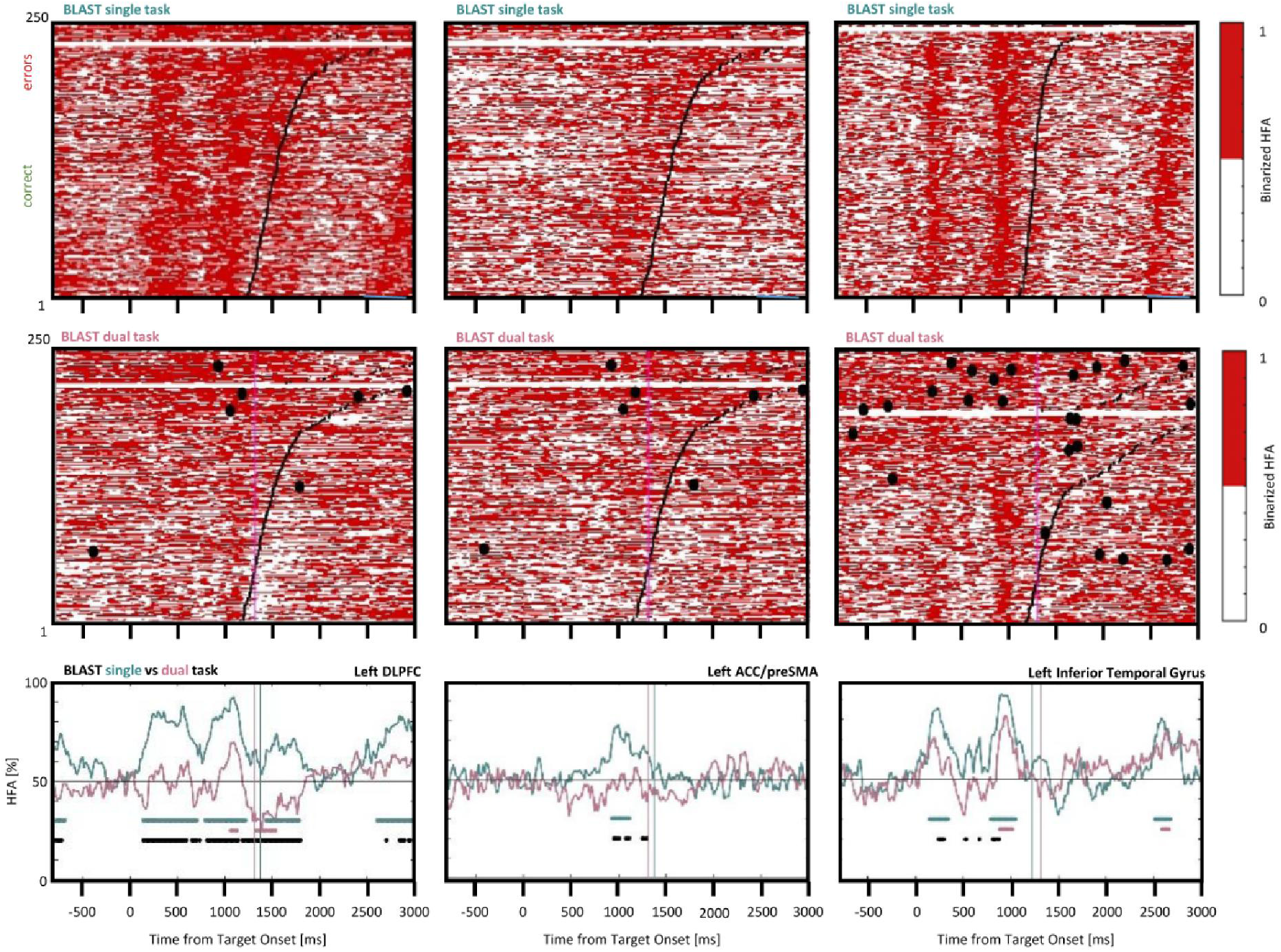
Interference control in the executive network and the language network. Comparison of BLAST-related activity in the ST and DT conditions for six sites (five of them supporting executive control in the anterior insula, DLPFC and ACC/preSMA, the last one supporting visual processes in the inferior temporal gyrus). For each site, BLAST-related neural activity is represented as a binarized matrix for the ST (top) and DT (middle) conditions. The curves at the bottom correspond to the average of those matrixes for the M fastest correct trials, where M equals half the number of correct trials in the DT condition. Horizontal lines indicate samples with a significant difference between the DT and ST curves (black), or a significant increase relative to the [-200:0 ms] pre-Target baseline, for the ST and DT conditions respectively (in green and red) (Kruskal-Wallis test, with a False-Discovery Rate correction for multiple comparisons). From left to right and top to bottom, iEEG sites and [Patients id] are: X’9 [P3]; X7 [P3]; E4 [P7]; K’10 [P5]; Z’2 [P5]; E’7 [P4].

## DISCUSSION

Our primary objective was to understand why retrieving information from long-term memory interferes with tasks, which require quasi-continuous attentive processing of visual stimuli. This type of interference is found in a large-class of real-life DT situations, such as driving while trying to remember a movie name during a ‘deep’ conversation (i.e. not a mere succession of speech automatisms). It is characteristic of all many situations, which combine the search for exogenous and endogenous information.

We specifically asked participants to search for information in long-term memory while processing frequent visual stimuli to produce an appropriate motor response, as during driving. That specific combination – BLAST and a Verbal Fluency Task – minimized interference at the sensory and motor level while preserving ecological validity with direct interpersonal interaction and self-scheduling of the memory retrieval, as during a normal conversation.

### Using iEEG to relax time-constraints and preserve ecological validity

Doing so, we admittedly departed from one of the founding principles of cognitive neuroscience: carefully controlled experiments, tailor-made to test contradicting predictions of existing theories (the “theory-driven” approach). But the choice was deliberate and characteristic of the growing field of naturalistic neuroscience (Sonkusare et al, 2019), which often starts from a real-life phenomenon and seeks an explanation from data collected in conditions preserving key features of the situation where it occurs (“situation-driven” approach) (e.g. Alm & Nilsson, 1994).

Precisely, one key feature of daily-life DT situations is that individuals are rarely pressed to perform both tasks fast and accurately: a driver is not forced to answer his passenger’s questions as fast as possible. In fact, most of us are sufficiently aware of the difficulty of multi-tasking to avoid any risky situation that imposes a double time constraint, unless the tasks have been widely practiced. Yet, most DT laboratory experiments combine two speeded-choice tasks, for a good reason: to control the relative timing of the two tasks and their sub-processes – from sensory analysis to response execution. This is exquisitely achieved by the Psychological Refractory Period (PRP) paradigm (Welford, 1952; Pashler, 1994), which sets the relative onset of the two tasks with millisecond precision. Nevertheless, the proper interpretation of PRP results requires that participants have genuinely tried to respond as fast and accurately as possible, and did not settle into a more comfortable pace to minimize DT interference. In contrast, applied research studies often allow such self-scheduling to retain ecological validity, but their primary goal is not so much to model the interference at a mechanistic level, but rather to assess the conditions that maximize the performance decrement and how participants manage the two tasks (Steinborn & Huestegge, 2017).

We predicted that intracranial EEG – as a time-resolved measure of task-induced brain activity – could provide a way to study the origin of the DT cost in naturalistic conditions despite reduced control over task timing. Given that our primary task – BLAST – had a very clear spatio-temporal neural signature visible at the single-trial level in iEEG recordings (Petton et al, 2019), it seemed possible to detect the impact of the VFT on individual components of BLAST – its putative « stages » – Identified from direct measurements rather than from a priori models of the task. The temporal precision of iEEG was also crucial to overcome one particular challenge of long-term memory retrieval: knowing when participants engage in the task. With iEEG, we could identify precise neural populations with abnormal high activity before the overt verbal responses, which is the only period when participants were surely involved in VFT memory retrieval. Our approach echoes previous studies that used fMRI and scalp EEG to combine their spatial and temporal precision (e.g. Sigman & Dehaene, 2008), but it bypasses the extreme difficulty of relating the sources of the two signals. In addition, the signal-to-noise ratio of scalp EEG – especially in the gamma frequency range (Yuval-Greenberg et al, 2008) – is not sufficient to measure the duration of task-related neural processes in single-trials, which limits the ability to relate EEG to behavioral variables.

### Dynamical incompatibility

We could show that the DT cost can be understood as a deviation from an optimal spatio-temporal dynamics, due to neuroelectric interferences in the cortical network supporting the primary task, BLAST. Interferences were observed throughout the broad, domain-general, Multiple-Demand Network (MDN) (Cole & Schneider, 2007; Rushworth, 2005; Dux et al, 2009; Crittenden & Duncan, 2014), including the DLPFC (the Inferior Frontal Sulcus and the Middle Frontal Gyrus), the Anterior Insula, the ACC/preSMA, and the Precentral Gyrus, as well as in process-specific areas: in the language network and high-level visual areas such as the Visual Word Form Area or the Fusiform Face Area, respectively. The temporal precision of iEEG revealed a situation of ‘dynamical incompatibility” with the VFT: a concept that we introduce to explain why two tasks can be mutually exclusive in naturalistic conditions despite the theoretical possibility of multiplexing. One might wonder for instance, why a phone conversation impairs driving performance while drivers can sample traffic information only once every second on average with no performance decrement (Han & Marois, 2013). In theory, such gaps might leave sufficient time for the cognitive processes of a normal conversation. During BLAST, we showed that although the task did not activate neural populations continuously, the temporal gaps were too short to nest VFT-related processes. This criterion is the defining feature of a “dynamical incompatibility” between the two tasks, which formally prevents time-sharing. Whether two tasks are dynamically compatible or not depends on the duration of the neural processes supporting those tasks, and the frequency at which they operate. It follows that the incompatibility can only be demonstrated via a detailed description of the subsecond dynamics of the two tasks and we believe, for this reason, that our iEEG study provides the first example in humans.

### The DT cost as a neuroelectric interference

Altogether, our data show that during BLAST, a secondary task can trigger strong and sustained responses in the gamma band in neural populations, which normally structure a precise temporal organization of activations/deactivations when performance is optimal. This effect is in itself a sufficient explanation of the poor BLAST performance in the DT condition, and by extrapolation, a plausible reason why driving and deep conversation do not mix well: for decades, clinicians have used trains of low amplitude electrical stimulations – most typically at 50 or 60 Hz, that is in the gamma range – to disrupt cognitive and motor functions, such as speech when stimulating specific sites in the left inferior frontal gyrus. The effects are so reliable that the procedure is considered as the clinical gold standard for presurgical functional brain mapping (Penfield & Rasmussen, 1950; Ojemann et al, 1989; Chassagnon et al, 2008). One possible explanation of the DT cost, therefore, is that the neuroelectric interferences caused by the secondary task in the gamma frequency range might mimic the effect of focal electrical stimulations. The advantage of that interpretation is that it creates a potentially fruitful bridge with a rich literature on the effect of electromagnetic stimulations (Direct Transcranial Current Stimulation, Transcranial Magnetic Stimulation) on task-performance (Hallett, 2007; Nitsche et al, 2008).

Our conclusions go beyond what a classic investigation of our paradigm, using behavioral measures and fMRI, could have revealed. We described the DT interference as a deviation from an optimal neural dynamics of the primary task, with little reference to well-identified « stages », which might seem like an impoverished explanation. Yet, if the objective is to understand why two tasks interfere with each other, the usual practice of decomposing tasks into successive stages is not an obligation and might in fact be misleading (Koch et al, 2018). For instance, our observations indicate clearly that during BLAST, perceptual analysis (in visual areas of the inferior temporal gyrus) and response selection (in the prefrontal cortex) largely occur in parallel while meaningful information is extracted from the letter array. As emphasized already by Jiang & Kanwisher (2003), and by Tombu and colleagues (2011) in their unified bottleneck model, peripheral and central processes should not be thought as necessarily separate and, we add, consecutive. This observation should alert us against the use of pre-conceived task models to explain DT interference from behavioral measures alone.

### A slower response selection process

In fact, many electrophysiological studies have seriously challenged the notion of sequential stages and the distinction between peripheral (sensory) analysis and central processes (response selection) (Gold & Shadlen, 2007; Hanks & Summerfield, 2017). This cornerstone of the original bottleneck theory is hard to reconcile with current models of perceptual decision-making, backed up by electrophysiological recordings in non-human primates (e.g. (Shadlen & Newsome, 2001). Rather than successive stages, such models propose that sensory and decision-level cortical areas jointly extract meaningful information to accumulate evidence in favor of one of several options, until a threshold is reached and a decision is made (Ratcliff & McKoon, 2008). Evidence accumulation is likely to occur in any task that involves “active vision” (sustained attention to visual stimuli to extract information), including BLAST. One additional problem with models that dissociate peripheral sensory analysis and central response selection is that some processes, such as memory encoding, are hard to associate with either stage : if the response selection of Task 1 was actually delayed by Task 2, were would sensory information be kept until the wait is over ? A waiting period, as postulated by the central bottleneck hypothesis, implies that sensory information is encoded and maintained into memory until the response selection component is free to process it. Yet memory encoding and maintenance also involve central components (in the prefrontal cortex), which complicates the design of a bottleneck theory.

Taken together, our data seem hardly compatible with any version of the bottleneck theory. The incompatibility was especially clear in the two patients with the most extensive iEEG sampling of the BLAST network: both displayed strong and sustained BLAST activations in sites affected and unaffected by the VFT. Their shared time course of activation was characteristic of a response selection process (sustained from the stimulus to the motor response).Importantly, activations were clearly visible in single-trials in both ST and DT conditions and, for sites free of VFT interferences, even for the slow or incorrect trials. In those recordings, the BLAST response was lengthened but not postponed, contrary to the prediction of the bottleneck theory. Therefore, the neuroelectric interference in the DT condition seemed to affect primarily the duration of a response selection component and its distributed neural counterpart. This provides another example of the benefit of iEEG, as a longer neural response would have been difficult to differentiate from a stronger one, with fMRI (Lachaux et al, 2007). This should warn against a simplistic interpretation of fMRI data from DT experiments. We see how in our experiment, it was particularly important to relate the timing of the neural response to major behavioral events (stimulus delivery, motor response, verbal responses to the VFT) in each recording site, to reveal anatomo-functional relationships. Although time-resolved fMRI has been used to show later activation peaks in the DT condition, compared to ST, in support of the bottleneck queue effect, we show here that only time-resolved recordings with millisecond precision at the single-trial level can show whether the response is actually postponed, or simply lengthened.

Our results are more easily interpreted within the resource-sharing theory, but with a line of reasoning sharply different from most fMRI studies. In most cases, fMRI leads to models of the DT cost based on three values per voxels: the BOLD signal when each task is performed in a ST condition (V1 and V2), and in the DT condition (V12). Although iEEG has undeniable limits (sparse cortical sampling, participants with a major neurological disorders, which can be dealt with following proper guidelines (Lachaux et al, 2012)), it provides for each recording site and condition the full time-course of a well-identified neural population. In fMRI, under additive summation in fMRI (when V12 < V1 + V2) is often taken as a strong indication that resources are shared between the two tasks (Just et al, 2001), but with the implicit assumptions of a linear relationship between resource attribution and the BOLD signal throughout the brain, and that more resources should lead to better performance. Comparatively, the iEEG observation that the neural activation is longer when reaction time is slower becomes problematic. Additionally, it has long been pointed out that fMRI cannot distinguish between the local recruitment of a single or two neighbor neural populations, and their serial or parallel activation, which limits the mechanistic interpretation of fMRI data during DT (Jiang & Kanwisher, 2003).

Our main evidence in favor of a resource-sharing account is that the neural activation was longer for the bad trials of BLAST, in many nodes of its network. As said earlier, BLAST involves a process of evidence accumulation, which can take longer in a dualtask condition. Kadohisa and colleagues showed that frontal and parietal neurons of monkeys trained to search for a target in a display develop a sensitivity for that target, and might therefore serve as detectors that receive information from visual areas during the accumulation process (Kadohisa et al, 2019). Yet, the sensitivity is reduced in the DT condition, suggesting that it might take longer for a perceptual decision to be made, if that decision requires that a given number of less selective neurons crossing a specific threshold. In that scenario, the long sought-after “resources” would correspond to the number of neurons optimally tuned to react to task-relevant stimuli in the primary task, exactly as proposed by Watanabe & Funahashi (2018). Interestingly, the duration of the BLAST response was also variable in the ST condition, which suggest that participants might spontaneously engage in other tasks, such as mind wandering, which might also slow down the evidence accumulation process.

### A Flexibility perspective on Dual-Tasking

In a recent review, Koch and colleagues emphasized the need for a flexibility perspective on multi-tasking, which emphasizes that participants can modify their strategy to adjust to momentary constraints, such as the obligation to perform a secondary task (Koch et al, 2018). Our data provide full support to that idea, showing how flexibility can even lead to radical changes in how the task is performed. During BLAST for instance, the target can be encoded, and to some degree searched for, using either verbal or visuo-spatial working memory (rehearsing the letter with covert speech or taking a visual ‘snapshot’). In some participants, whose strategy could be deduced from the activity of regions supporting phonological processes – such as the lower Precentral Gyrus – our data revealed striking instances where those regions withdrew from the BLAST task for the most successful trials in the DT session. Considering the fact that the VFT heavily relies on verbal processes (Shao et al, 2014) it seems like an efficient way of coping with that specific combination of tasks. Interestingly, that effect was not observed for the least successful trials, suggesting that the participants might have found it difficult to stabilize such an alternate and possibly less natural strategy.

Yet, most authors would agree that flexibility has some limits, and that regions which participate in all tasks that require cognitive control – within the Multiple-Demand Network (Duncan & Owen, 2000; Duncan, 2010); should create unavoidable interferences, unless tasks are extensively practiced. We were therefore utterly surprised to observe that some participants found a way, in the DT condition, to reduce the response of the ACC and the AI which constitute together the core of the executive Multi-Demand Network (core eMDN), in charge of “executive processing and cognitive control by initiating and maintaining cognitive sets, coordination behavioral responses and guiding behavior in general” (Shashidhara et al, 2019). BLAST is a demanding task designed to prevent automatization (Borkowski et al, 1967) and it logically activates the core eMDN consistently, at least in the ST condition. Therefore, we certainly did not expect that BLAST could be performed efficiently by some participants with virtually no activation of the core eMDN, especially during the best trials of the DT session, when more control should be needed. One might argue that the role of the AI/ACC extends to error monitoring (Dosenbach et al, 2007; Duncan, 2001; Klein et al, 2013) which could explain their strong activation during the worst trials of BLAST in the DT condition. However, that suggestion is incompatible with our observation that those two regions were systematically active in the ST condition, even for successful trials. One remaining explanation is that participants might have automatized BLAST during the ST session, and therefore exerted less executive control during the DT session, which came after the ST session. However, BLAST is so long and repetitive, with 250 identical trials, that signs of automatization should have been already visible in the ST condition when comparing AI/ACC activity between the beginning and the end of the experiment. This was not the case.

We believe that our data rather support the following scenario: in the DT condition and consciously or not, participants tend to exert cognitive control on the difficult VFT rather than on BLAST, with surprisingly little impact on BLAST performance. It follows that BLAST can be performed fast and accurately with less control (i.e. less activation of the eMDN), which provides an unexpected strategy to reduce DT interference: performing one of the two tasks “as if” it was fully automatized.

Since cognitive control has been associated with mental fatigue, one might then wonder why participants did not use that strategy also in the ST condition. One possible explanation is that the need for control is so deeply rooted that one must be forced to ‘let go’ by a secondary task, when control cannot be exerted upon both equally. The implication of that finding is rather profound, as it suggests that we might be exerting excessive cognitive control, and therefore excessive mental effort, during tasks that could be done almost effortlessly. This conclusion is reminiscent of a study that used the attentional blink phenomenon to show that control participants, compared to meditation experts, over-process task-relevant stimuli (Han et al, 2019) — excessive effort, again. More generally, it should remind us that focused attention should be dissociated from mental effort, as clearly demonstrated by multiple insightful contributions in a seminal book on that topic (Slagter et al, 2007)

## CONCLUSION

To conclude, we showed that the neural basis of the DT cost can be studied in partially unconstrained naturalistic conditions, provided time-resolved measures of the large-scale neural dynamics underlying the two tasks. iEEG provides two levels of explanation of the DT interference : a rather unsophisticated and yet simple explanation is that the secondary task generates a neuroelectric interference in the gamma range which might mimic the effect of direct cortical electrical stimulations at 50 or 60 Hz and trigger a deviation from the optimal large-scale spatio-temporal dynamics of the primary task. Yet, our data are also consistent with a second, non-exclusive, explanation where response selection is slowed down by the secondary task because neurons involved in the perceptual decision are less optimally tuned to the main task (Kadohisa et al, 2019). Only single-neurons recordings in humans could demonstrate that proposal, already validated in non-human primates.

This effect can be framed within the resource theory framework, but is harder to reconcile with the bottleneck model. Yet, one should be aware that our current understanding of DT should not be limited by the historical metaphors of the field, nor by the experimental measures accessible so far – mostly behavioral and BOLD data. Many initial models of the DT cost, based solely on behavioral data and smart experimental manipulations, claimed strong explanatory power with virtually no brain implementation. Then, a wealth of neuroimaging studies led to neurocognitive models of DT, with increased explanatory power, but still missing one essential ingredient: time. Because fMRI lacked fine temporal precision, the interference between tasks could not be thought as an interference between optimal spatio-temporal dynamics, with key dynamical concepts such as duration, relative timing, simultaneity, propagation and dynamical incompatibility. Hopefully, the time is now ripe for local electrophysiology to become a major avenue of DT research in the near future, as rightfully proposed by Watanabe & Funahashi (2018).

## DATA AND CODE AVAILABILITY STATEMENT

The data that support the findings of this study are available from the corresponding author, JP. L., upon reasonable request.

