## Supplemental Table 1 for "THE DUAL-TASK COST IS DUE TO NEURAL INTERFERENCES DISRUPTING THE OPTIMAL SPATIO-TEMPORAL DYNAMICS OF THE COMPETING TASKS"

### TABLES

Table I

| **Patient ID** | **iEEG site** | **X** | **Y** | **Z** | **Freesurfer labelization** |
| --- | --- | --- | --- | --- | --- |
| P1 | R9 | 67 | 0 | 10 | Subcentral gyrus |
| P1 | F6 | 41 | -50 | -13 | Lateral Occipito-Temporal Sulcus |
| P2 | M9 | 38 | 8 | 33 | Inferior Frontal Sulcus |
| P3 | X7 | 34 | 25 | 2 | Anterior Insula |
| P3 | X’9 | -30 | 23 | 5 | Anterior Insula |
| P4 | R’9 | -61 | 7 | 16 | Precentral Gyrus |
| P4 | G’13 | -45 | 33 | 21 | Inferior Frontal Sulcus |
| P4 | F’9 | -28 | 44 | 30 | Middle Frontal Sulcus |
| P4 | Y’13 | -31 | 50 | 27 | Middle Frontal Gyrus |
| P4 | E’7 | -50 | -37 | -21 | Inferior Temporal Gyrus |
| P5 | S’9 | -41 | 0 | 40 | Precentral Sulcus |
| P5 | K’10 | -42 | 35 | 12 | Inferior Frontal Sulcus |
| P5 | Z’2 | -11 | 19 | 46 | Anterior Cingulate Gyrus/Sulcus |
| P5 | L’3 | -36 | -52 | -11 | Lateral Occipito-Temporal Sulcus |
| P6 | D’7 | -55 | -42 | 8 | Superior Temporal Sulcus |
| P7 | E4 | 41 | 17 | 4 | Anterior Insula |
| P7 | S2 | 11 | 16 | 52 | Anterior Cingulate Gyrus/Sulcus |
| P9 | G’12 | -45 | 41 | 14 | Middle Frontal Gyrus |
| P11 | Q’2 | -40 | 25 | 14 | Inferior Frontal Sulcus |
| P11 | X5 | 30 | 20 | -6 | Anterior Insula |
| P11 | X’9 | -30 | 23 | 5 | Anterior Insula |
| P11 | C’12 | -60 | -37 | -12 | Middle Temporal Gyrus |
| P12 | X7 | 33 | 17 | 10 | Anterior Insula |
| P12 | A9 | 52 | -3 | -20 | Middle Temporal Gyrus |

Table I. MNI-coordinates for each site displayed in the figures with the anatomical labelization provided by the Freesurfer software (corrected when obviously inaccurate) (https://surfer.nmr.mgh.harvard.edu).
